# Role of mucin gene for growth in Anas platyrynchos - a novel report

**DOI:** 10.1101/2022.01.29.478287

**Authors:** Anuj kumar Murmu, Aruna Pal, Manti Debnath, Argha Chakraborty, Subhomoy Pal, Samiddha Banerjee, Abantika Pal, Nilotpal Ghosh, Utpal Karmakar, Rajarshi Samanta

**Author notes:** These authors have contributed equally (Anuj kumar Murmu, Aruna Pal, Manti Debnath). Corresponding Author: Assistant Professor (Animal Genetics and Breeding). West Bengal University of Animal and Fishery Sciences.37, K.B.Sarani, Kolkata-37, West Bengal, India. E mail.

## Abstract

Mucin gene is expressed at the mucous membrane of the inner layer of the internal organs. Intestinal mucin 2 (MUC2), a major gel-forming mucin, represents a primary barrier component of mucus layers. This is the first report of role of mucin gene in growth traits in animals. In the current study, we had randomly studied Bengal ducks (Anas platyrynchos) reared from day old to 10 weeks of age under organized farm and studied the growth parameters as well as body weight and average daily body weight gain. We had characterized mucin gene for Bengal duck and observed glycosylation and EGF1 (EGF like domain signature) as important domain for growth traits in duck. We observed better expression profile for mucin gene in high growing ducks in comparison to that of lower growing ducks with real time PCR. Hence mucin gene may be employed as a marker for growth traits.

## Introduction

Mucus is a viscous gel-like material covering the gastro-intestinal mucosal surface. The entire surface of the chicken gastrointestinal tract is covered by a layer of mucus that functions as a diffusive barrier between the intestinal lumen and absorptive cells. The mucins are the main component of the mucus layer, which are produced and secreted by goblet cells. Mucins are high-molecular weight glycoproteins (50–80% O-linked oligosaccharides) produced by epithelial tissues in most animals. These glycoproteins are found in mucus (e.g. saliva, gastric juice, etc.) and secreted by mucous membranes to lubricate or protect body surfaces of vertebrates and they have a central role in maintaining epithelial homeostasis (Marín *et al*., 2012). The mucus layer is part of the innate host response, protecting against luminal microflora, preventing gastrointestinal pathologies, and participating in the processes of nutrient digestion and absorption (Forstner *et al*., 1995). A decrease in mucin synthesis in poultry could compromise the mucus layer and reduce nutrient utilization (Horn *et al*., 2009).

Mucins are divided into secretory and membrane-bound types, based on their forms. Membrane-bound mucins (MUC1, MUC3A, MUC3B, MUC4, MUC12, MUC13, MUC15, MUC16, MUC17, MUC20, and MUC21) exhibit hydrophobic sequences or “transmembrane domains” responsible for anchoring them in the lipid bilayer and have C-terminal peptides that enter the cytosol. The secretory mucins (MUC2, MUC5AC, MUC5B, MUC6, MUC8, and MUC19) with one exception (MUC7) possess one or several von Willebrand factor (vWF)-like D domains, and cysteine-rich peptides, which function in the oligomerization of mucin monomers and in packaging into secretory vesicles (Lehmann et al., 1989; Perez-Vilar and Hill, 1999; Moniaux et al., 2001; Chen et al., 2004; Higuchi et al., 2004; Itoh et al., 2008;). According to whether they are capable of forming a gel, the secretory mucins can be further divided into two subtypes, gel-forming and soluble MUCs. Interestingly, MUC2, MUC5AC, MUC5B, and MUC6 belong to gel-forming MUCs and are located near each other on human chromosome 11p15.5 (Pigny et al., 1996).

Mucin is the major constituent of the mucus layer and serves a crucial role in protecting the gut from acidic chyme, digestive enzymes, and pathogens. In addition to its protective functions, mucin is involved in filtering nutrients in the gastrointestinal tract (GIT) and can influence nutrient digestion and absorption (Montagne *et al*., 2004). Any component, dietary or environmental, that induces changes in mucin dynamics has the potential to affect viscosity, integrity of the mucus layer, and nutrient absorption. Mucins in general contain many threonine and serine residues, which are extensively O-glycosylated. Due to this profound glycosylation, mucins have a filamentous conformation.

Ducks were observed to be mostly forager with better disease resistance (Pal et al., 2021), higher adaptability and hardy breed (Pal, A. 2020, Pal et al, 2019). Apart from metagenomic studies, gut of the foraging birds form a major role in providing immunity, and hardy nature of the birds, particularly ducks raised under semi-intensive system of management. Although reports were available for role of mucin in nutrient absorbtion and digestion, no report as such was available for the role of mucin gene in growth traits at molecular level. Growth trait is important for both meat and egg production of the duck. We had earlier studied the importance of certain genes in growth for livestock (Pal, A. 2014, Pal et al., 2020, Pal et al., 2010, Pal et al., 2012, Pal et al., 2020, Pal et al., 2017). Growth is a polygenic quantitative trait, hence it is important to study certain genes, like mucin gene, affecting gut integrity and nutrient absorption senario. In our lab, we are studying growth parameters including body weight and growth parameters for livestock species (**19-21** Pal et al. 2010, Pal, A, 2020, Pal et al., 2012, Pal et al., 2004, Pal et al., 2017, Pal et al., 2006, Pal et al., 2020, Pal et al., 2019). Hence the current study was designed to assess the role of mucin gene for growth of ducks through differential mRNA expression profiling.

## Materials and Methods

### Birds

The present study was conducted on Bengal ducks under organized farming system. The present investigation was carried out at livestock farm of West Bengal University of Animal and Fishery Sciences, Belgachia, Kolkata, West Bengal. The farm data and samples from ducks were collected from March to May, 2020. Later samples were processed for appropriate molecular biology work. The annual mean temperature is 26.8 °C (80 °F); monthly mean temperatures range from 19 °C to 30 °C (67 °F to 86 °F) and maximum temperatures can often exceed 40 °C (104 °F) during May–June.

Sixty (n=60) numbers of healthy indigenous duckling (*Anas platyrhynchos*) were used for the present study. The ducklings were properly maintained under standard feeding and management. Duck eggs were collected from field and subjected to hatching for average 28 days and reared in our farm from day old stage. The ducklings (Day old to 2 weeks of age) were given starter ration containing 20% crude protein and 2750 Kcal ME per Kg. The first growing phases (3-8 weeks), received the feed composed of 18% crude protein and 2750 Kcal ME and the second growing phases (9-20 weeks), received the feed composed of 15% crude protein and 2700 Kcal ME per Kg (**22** NRC,1994). Studies were conducted with ethical approvals obtained from Institutional Animal Ethics Committee, West Bengal University of Animal and Fishery Sciences.

### Data and Sample collection

At the end of each week, the ducks were weighed using digital weighing balance to determine the body weight in gram. Other morphometric parameters i.e., shank length, breast width, keel length, drumstick length, neck length, body length, chest girth and wing length was measured with measuring tape for 8 weeks and expressed in centimeter (cm). Daily body weight gain was assesed through computational approach.

Daily body weight gain= Body weight/age of the bird.

Based on the available data, the entire duckling population data into high body weight (better growing) as values above mean +standard deviation, and low body weight (less growing) as values less than mean –standard deviation. Trend for average daily body weight gain have been depicted. Similar groups were also observed for average daily body weight gain and biomorphometric traits of the ducks.

Duodenum and caecum were collected from the duck belonging to two groups-higher and lower body weights.

### Characterization of Mucin 2 gene with respect to growth parameters

All experiments were conducted in accordance with relevant guidelines and regulations of Institutional Animal Ethics committee and all experimental protocols were approved by the Institutional Biosafety Committee, West Bengal University of Animal and Fishery Sciences, Kolkata.

The total RNA was isolated from the duodenum and caecum of Duck by Trizol method (**23-31**) and was further used for cDNA synthesis.

### Materials

Taq DNA polymerase, 10X buffer, dNTP were purchased from Invitrogen, SYBR Green qPCR Master Mix (2X) was obtained from Thermo Fisher Scientific Inc. (PA, USA). L-Glutamine (Glutamax 100x) was purchased from Invitrogen corp., (Carlsbad, CA, USA). Penicillin-G and streptomycin were obtained from Amresco (Solon, OH, USA). Filters (Millex GV. 0.22 μm) were purchased from Millipore Pvt. Ltd., (Billerica, MA, USA). All other reagents were of analytical grade.

### Synthesis, Confirmation of cDNA and PCR Amplification of Mucin2 gene

The 20 □ μL reaction mixture contained 5 □ μg of total RNA, 0.5 □ μg of oligo dT primer (16–18 □ mer), 40□U of Ribonuclease inhibitor, 10□M of dNTP mix, 10 □ mM of DTT, and 5□U of MuMLV reverse transcriptase in reverse transcriptase buffer. The reaction mixture was gently mixed and incubated at 37°C for 1 hour. The reaction was stopped by heating the mixture at 70°C for 10 minutes and chilled on ice. The integrity of the cDNA was checked by PCR. To amplify the full-length open reading frame (ORF) of gene sequence, a specific primers pair was designed based on the mRNA sequences of *Gallus gallus* by DNASTAR software. The primers have been listed in Table 1. 25□μL reaction mixture contained 80-100□ng cDNA, 3.0 μL 10X PCR assay buffer, 0.5 □μL of 10 mM dNTP, 1□U Taq DNA polymerase, 60□ng of each primer (as in Table 1), and 2□mM MgCl2. PCR-reactions were carried out in a thermocycler (PTC-200, MJ Research, USA) with cycling conditions as, initial denaturation at 94°C for 3□min, denaturation at 94°C for 30□ sec, varying annealing temperature (as mentioned in Table 1) for 35□sec, and extension at 72°C for 3 □ min was carried out for 35 cycles followed by final extension at 72°C for 10□min.

**Table 1:**
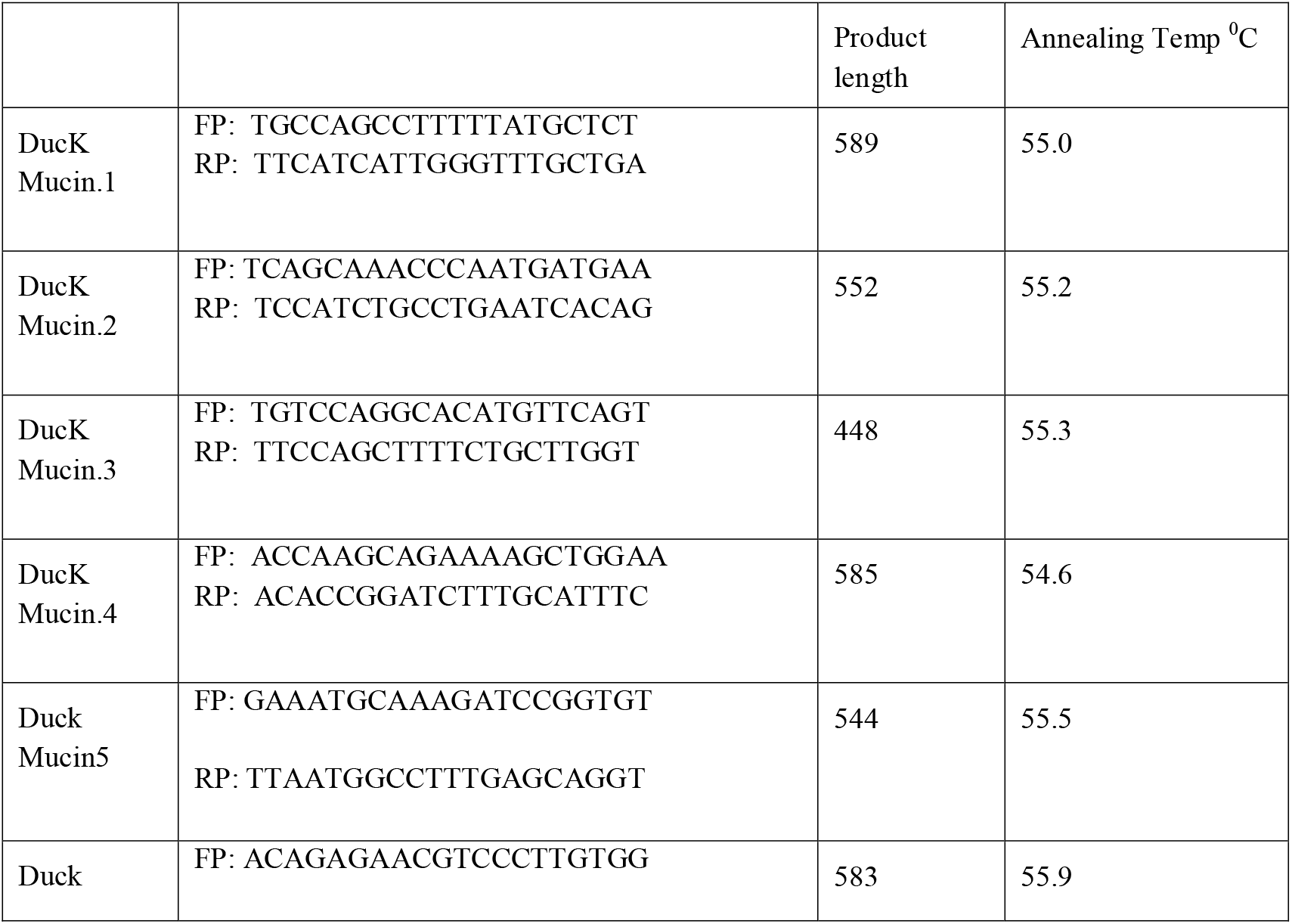
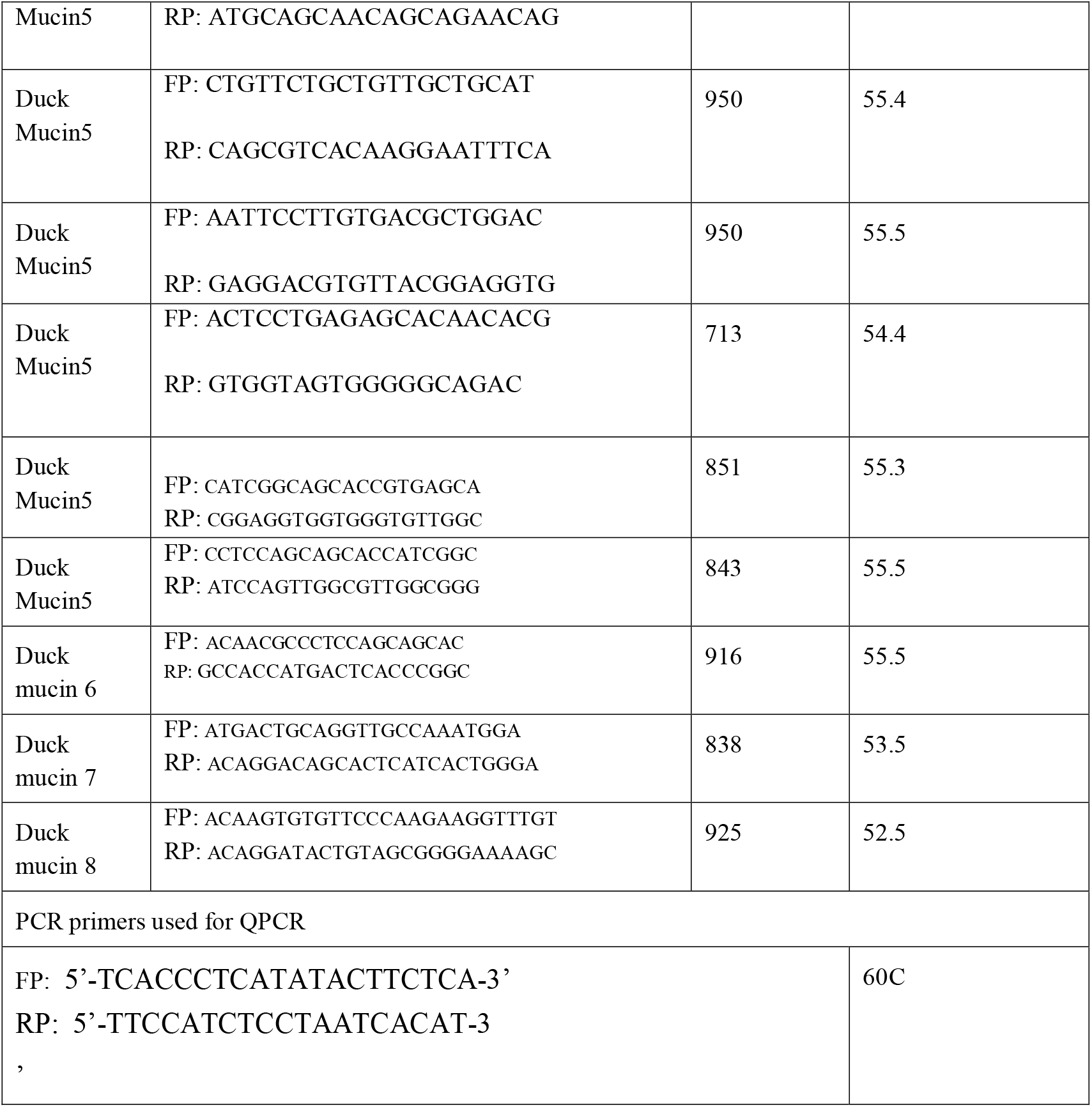
PCR primers for Mucin 2 in Duck:

**Table 1:** Body weight at successive weeks of life for two different levels of growth.

| Age | Group I (High body weight, g) | Group II (Low body weight, g) |
| --- | --- | --- |
| 1<sup>st</sup> week | 128.6± 3.331<sup>a</sup> | 53.0±1.176<sup>b</sup> |
| 2<sup>nd</sup> week | 195.478±5.751<sup>a</sup> | 118.667±1.153<sup>b</sup> |
| 3<sup>rd</sup> week | 373.3± 5.86<sup>a</sup> | 209.0± 1.0522<sup>b</sup> |
| 4<sup>th</sup> week | 529.375±7.074<sup>a</sup> | 275.0± 2.0558<sup>b</sup> |
| 5<sup>th</sup> week | 688.9± 10.907<sup>a</sup> | 406.571± 1.9236<sup>b</sup> |
| 6<sup>th</sup> week | 773.862± 11.091<sup>a</sup> | 454.0± 3.4616<sup>b</sup> |
| 7<sup>th</sup> week | 1035.6± 12.391<sup>a</sup> | 534.778± 3.4354<sup>b</sup> |
| 8<sup>th</sup> week | 1134.667± 13.924<sup>a</sup> | 585.778± 4.0664<sup>b</sup> |
| 9<sup>th</sup> week | 1246.0±13.488<sup>a</sup> | 588.0± 4.4806<sup>b</sup> |
| 10<sup>th</sup> week | 1385.548± 14.245 <sup>a</sup> | 927.222± 7.259 <sup>b</sup> |

For all values the level of significance $P \leq 0.01$

### cDNA Cloning and Sequencing

PCR amplicons verified by 1% agarose gel electrophoresis were purified from gel using Gel extraction kit (Qiagen GmbH, Hilden, Germany) and ligated into a pGEM-T easy cloning vector (Promega, Madison, WI, USA) following manufacturers’ instructions. The 10 μL of the ligated product was directly added to 200□μL competent cells, and heat shock was given at 42°C for 45 □ sec. in a water bath, and cells were then immediately transferred on chilled ice for 5 □ min., and SOC was added. The bacterial culture was pelleted and plated on LB agar plate containing Ampicillin (100□mg.mL) added to agar plate @ 1□:□1000, IPTG (200□mg. mL) and X-Gal (20□mg. mL) for blue-white screening. Plasmid isolation from overnight-grown culture was done by small-scale alkaline lysis method. Recombinant plasmids were characterized by PCR using gene-specific primers and restriction enzyme digestion based on reported nucleotide sequence for chicken. The enzyme EcoRI (MBI Fermentas, USA) is used for fragment release. Gene fragment insert in the recombinant plasmid was sequenced by an automated sequencer (ABI prism) using the dideoxy chain termination method with T7 and SP6 primers (Chromous Biotech, Bangalore).

### Sequence Analysis

The nucleotide sequence so obtained was analyzed for protein translation, sequence alignments, and contigs comparisons by DNASTAR Version 4.0, Inc., USA. The novel sequence was submitted to the NCBI Genbank and accession number was obtained which is available in public domain now.

### Study of Predicted Mucin 2 peptide Using Bioinformatics Tools

The predicted peptide sequence of Mucin 2 of indigenous duck was derived by Edit sequence (Lasergene Software, DNASTAR) and then aligned with the peptide of other chicken breed and avian species using Megalign sequence Programme of Lasergene Software, DNASTAR (**32-34**). Prediction of the signal peptide of the Mucin 2 gene was conducted using the software (Signal P 3.0 Sewer-prediction results, Technical University of Denmark). Estimation of Leucine percentage was conducted through manually from the predicted peptide sequence. Di-sulfide bonds were predicted using suitable software (http://bioinformatics.bc.edu/clotelab/DiANNA/) and by homology search with other species.

Protein sequence-level analysis study was carried out with specific software (http://www.expasy.org./tools/blast/) for determination of leucine-rich repeats (LRR), leucine zipper, N-linked glycosylation sites, detection of Leucine-rich nuclear export signals (NES), and detection of the position of GPI anchor. Detection of Leucine-rich nuclear export signals (NES) was carried out with NetNES 1.1 Server, Technical University of Denmark. Analysis of O-linked glycosylation sites was carried out using NetOGlyc 4 server (http://www.expassy.org/), whereas the N-linked glycosylation site was detected by NetNGlyc 1.0 software (http://www.expassy.org/). Detection of Leucine-zipper was conducted through Expassy software, Technical University of Denmark^**30-34**^. Regions for alpha-helix and beta-sheet were predicted using NetSurfP-Protein Surface Accessibility and Secondary Structure Predictions, Technical University of Denmark^**30-34**^. Domain linker prediction was done according to the software developed^**30-35**^. LPS-binding site, as well as LPS-signaling sites^**37**^, were predicted based on homology studies with other species polypeptide.

### Three-dimensional structure prediction and Model quality assessment

The templates which possessed the highest sequence identity with our target template were identified by using PSI-BLAST (http://blast.ncbi.nlm.nih.gov/Blast). The homology modeling was used to build a 3D structure based on homologous template structures using PHYRE2 server^**38**^. The 3D structures were visualized by PyMOL (http://www.pymol.org/) which is an open-source molecular visualization tool. Subsequently, the mutant model was generated using PyMoL tool. The Swiss PDB Viewer was employed for controlling energy minimization. The structural evaluation along with a stereochemical quality assessment of predicted model was carried out by using the SAVES (Structural Analysis and Verification Server), which is an integrated server (http://nihserver.mbi.ucla.edu/SAVES/). The ProSA (Protein Structure Analysis) webserver (https://prosa.services.came.sbg.ac.at/prosa) was used for refinement and validation of protein structure^**39**^. The ProSA was used for checking model structural quality with potential errors and the program shows a plot of its residue energies and Z-scores which determine the overall quality of the model. The solvent accessibility surface area of the Mucin 2 gene was generated by using NetSurfP server (http://www.cbs.dtu.dk/services/NetSurfP/). It calculates relative surface accessibility, Z-fit score, the probability for Alpha-Helix, probability for beta-strand and coil score, etc. TM align software was used for the alignment of 3 D structure of IR protein for different species and RMSD estimation to assess the structural differentiation^**40**^. The I-mutant analysis was conducted for mutations detected to assess the thermodynamic stability^**41**^. Provean analysis was conducted to assess the deleterious nature of the mutant amino acid^**42**^. PDB structure for 3D structural prediction of Mucin 2 gene for duck was carried out through PHYRE software^**38**^. Protein-protein interaction have been studied through String analysis^**43,44**^.

### Real time PCR

Total RNA was estimated from duodenum and caecum of duck from high bodyweight and low body weight group by Trizol method and quantitative analysis of total RNA were performed using formaldehyde gel electrophoresis. The 28S rRNA and 18S rRNA were demarcated the quality of RNA.First strand cDNA was synthesised by the process of reverse transcriptase polymerase chain reaction (rt-PCR) in the automated temperature maintained thermocycler mechine. M-MLVRT (200 u/μl) was used as reverse transcriptase enzyme. All the primers were designed using primer 3 software (v. 0. 4.0) as per the recommended criteria. The primers used are listed in Table1. Equal amount of RNA (quantified by Qubit fluorometer, Invitrogen), wherever applicable, were used for cDNA preparation (Superscript III cDNA synthesis kit; Invitrogen). All qRT-PCR reactions were conducted on ABI 7500 fast system. Each reaction consisted of 2 μl cDNA template, 5 μl of 2X SYBR Green PCR Master Mix, 0.25 μl each of forward and reverse primers (10 pmol/μl) and nuclease free water for a final volume of 10 μl. Each sample was run in duplicate. Analysis of real-time PCR (qRT-PCR) was performed by delta-delta-Ct (ΔΔCt) method^**45**^.

The entire reactions were performed in triplicate (as per MIQE Guidelines) and experiment repeated twice, in 20μl reaction volume, using FastStart Essential DNA Green Master (Himedia) on ABI 7500 system.

### Statistical analysis

Descriptive statistics with mean and standard error were estimated through SYSTAT package for the expression level analyzed through real time PCR and presented accordingly in graph.

The model emloyed:

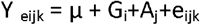

where Y _eijk_ = kth observation of the target trait, μ = Overall mean,

Gi = Fixed effect of ith genotype corresponding to expression level, Aj= Fixed effect of jth bird, e_ijk_= Random error

Expression level with real time PCR was estimated as 2^−ΔΔCt^

## Result

### Characterization of mucin gene in Duck

Mucin 2 coats the epithelia of the intestines, airways, and other mucus membrane-containing organs. Certain important domains for Mucin 2 protein has been detected. It was attempted to explore the growth potential ability of the domains. Signal peptide was detected at amino acid position 1-18 for the predicted 3D protein structure of Mucin 2 gene. Certain important domains have been identified as VWFD (VonWillebrand factor) domain at amino acid sites of 33-236 (red sphere), 388-601 (forest green), 859-1065 (orange sphere), 2945-3164 (**Fig 1**). Other identified domains were VWFC-1 at sites of amino acid 3293-3343, 3402-3448. VWFC-2 domains were predicted at amino acid location of 3275-3344, 3382-3449. Other two important sites were CTCK1 (C terminal cystine knot signature), CTCK2 (C terminal cysteine knot domain profile).The sites for domain of CTCK1 are 3580-3618 and CTCK2 at aa position 3532-3619. Sites for disulphide bonds were detected at aa positions 56-64, 411-419, 882-890, 2968-2976, 3532-3581, 3557-3611, 3561-3613 (**Fig 2**).

**Fig 1:**
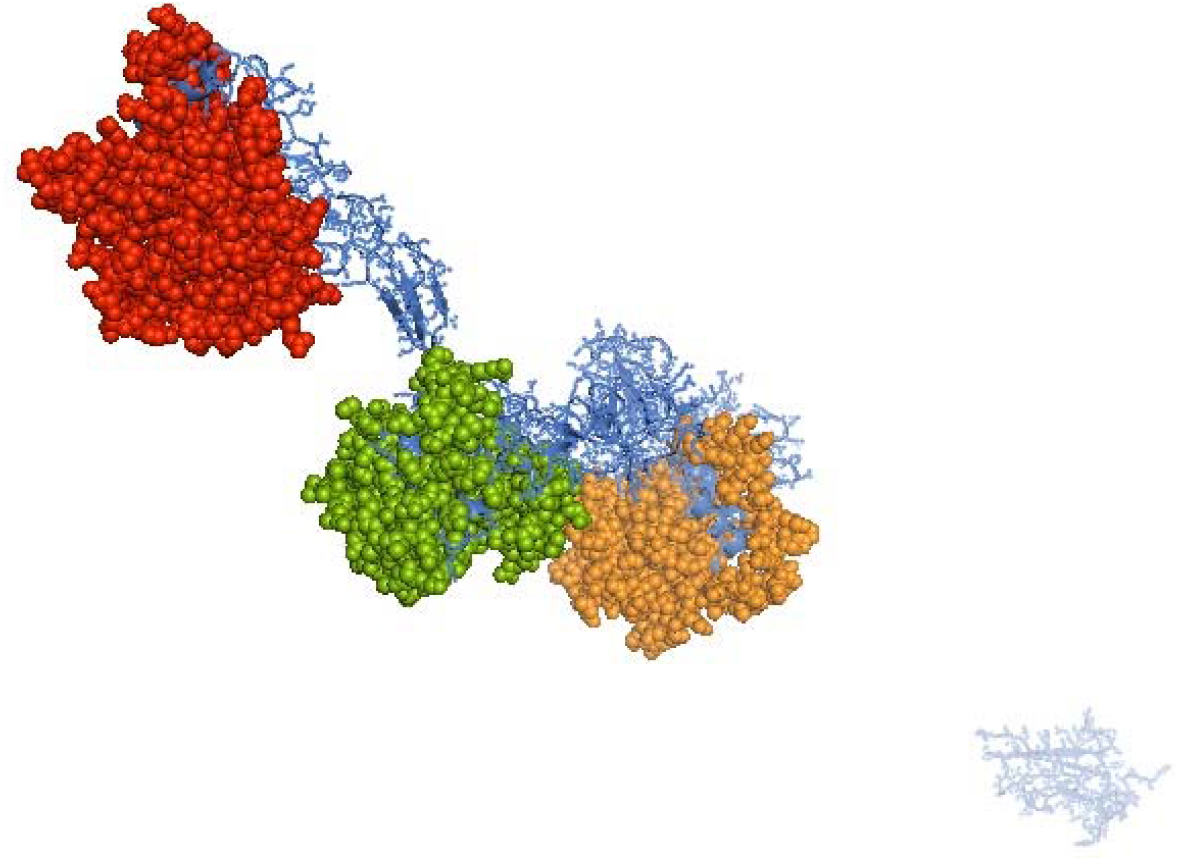
Domain for Mucin2-VWFD (VonWillebrand factor) domain at amino acid sites of 33-236, 388-601, 859-1065, 2945-3164 for the mucin 2 gene of duck.

**Fig 2:**
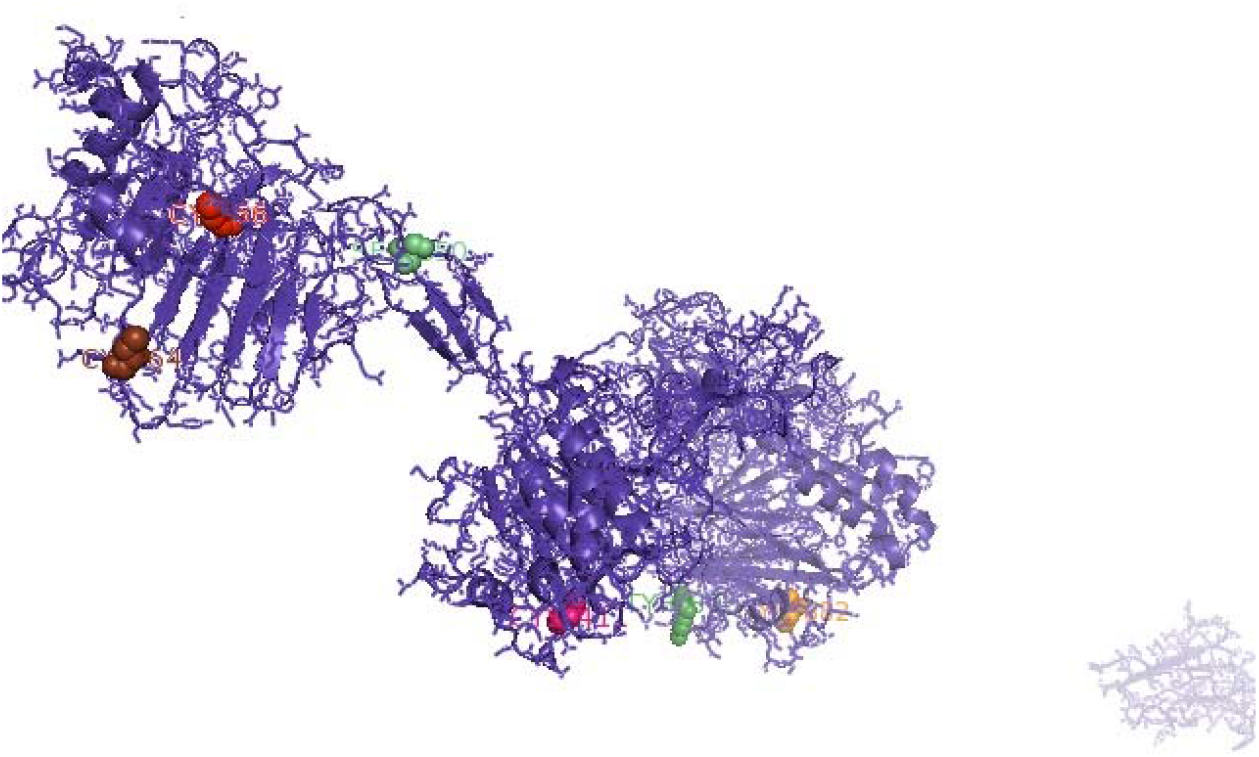
Sites for disulphide bonds were detected at aa positions 56-64, 411-419, 882-890, 2968-2976, 3532-3581, 3557-3611, 3561-3613 for the mucin 2 gene of duck

In this current context, we are concerned with growth parameters for mucin gene. We had identified an important domain involved with growth as EGF1 (EGF like domain signature) at sites of 3296-3307. Certain important observations were revealed. Since disulphide bonds were not included in this region, it is presumed to be open. EGF1 domain is inserted within VWFC-1 and VWFC-2.

The sites for glycosylation is very important. The sites for N-linked glycosylation and O-linked glycosylation have been listed in Supplementary Table. 22 sites for N-linked glycosylation and sites for O linked glycosylation has been predicted.

The pictorial description for secondary structure for Mucin 2 gene is being described in **Fig 3**.It represents the site for alpha helix, beta sheet and loop structure for the mucin gene.

**Fig 3:**
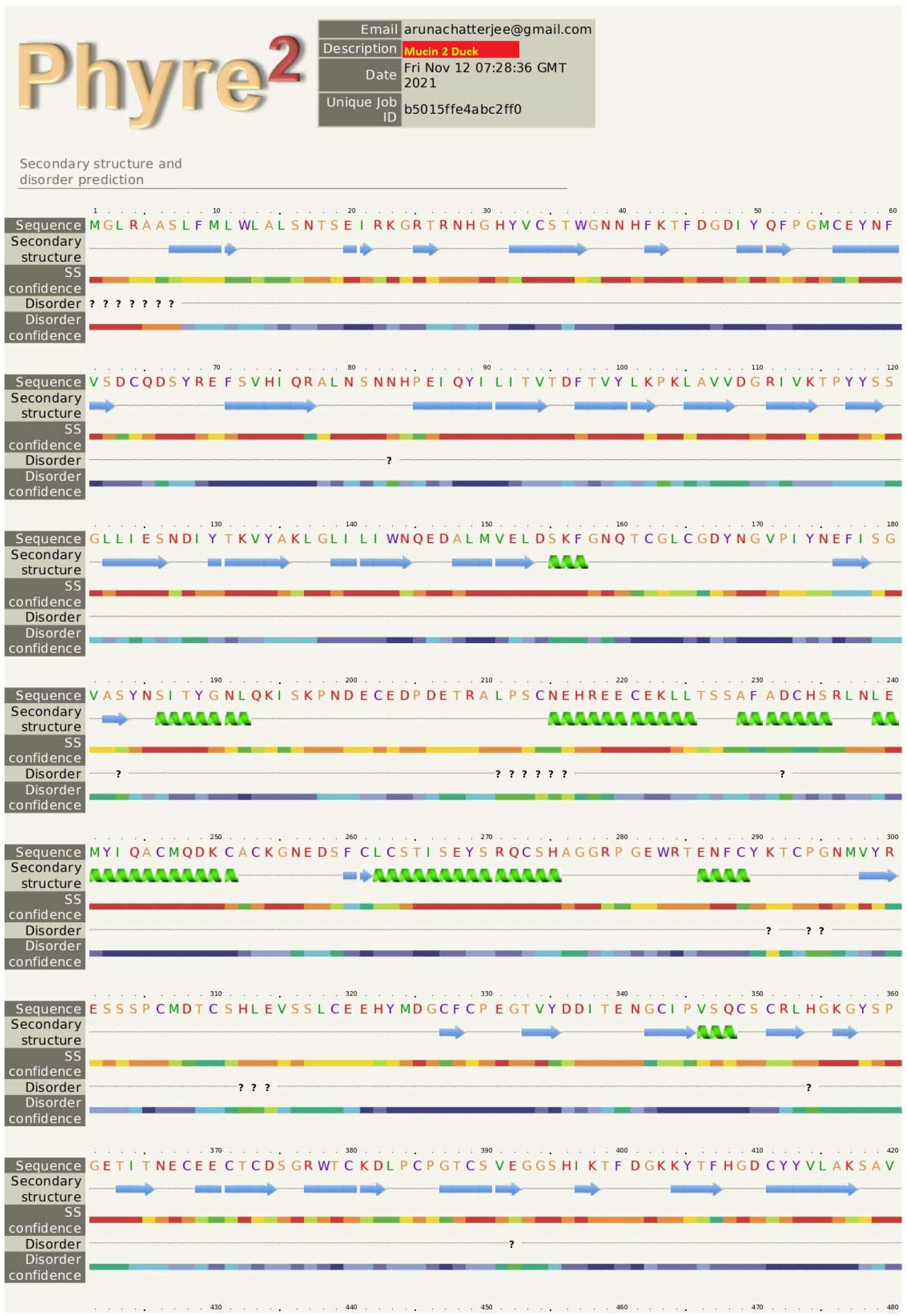
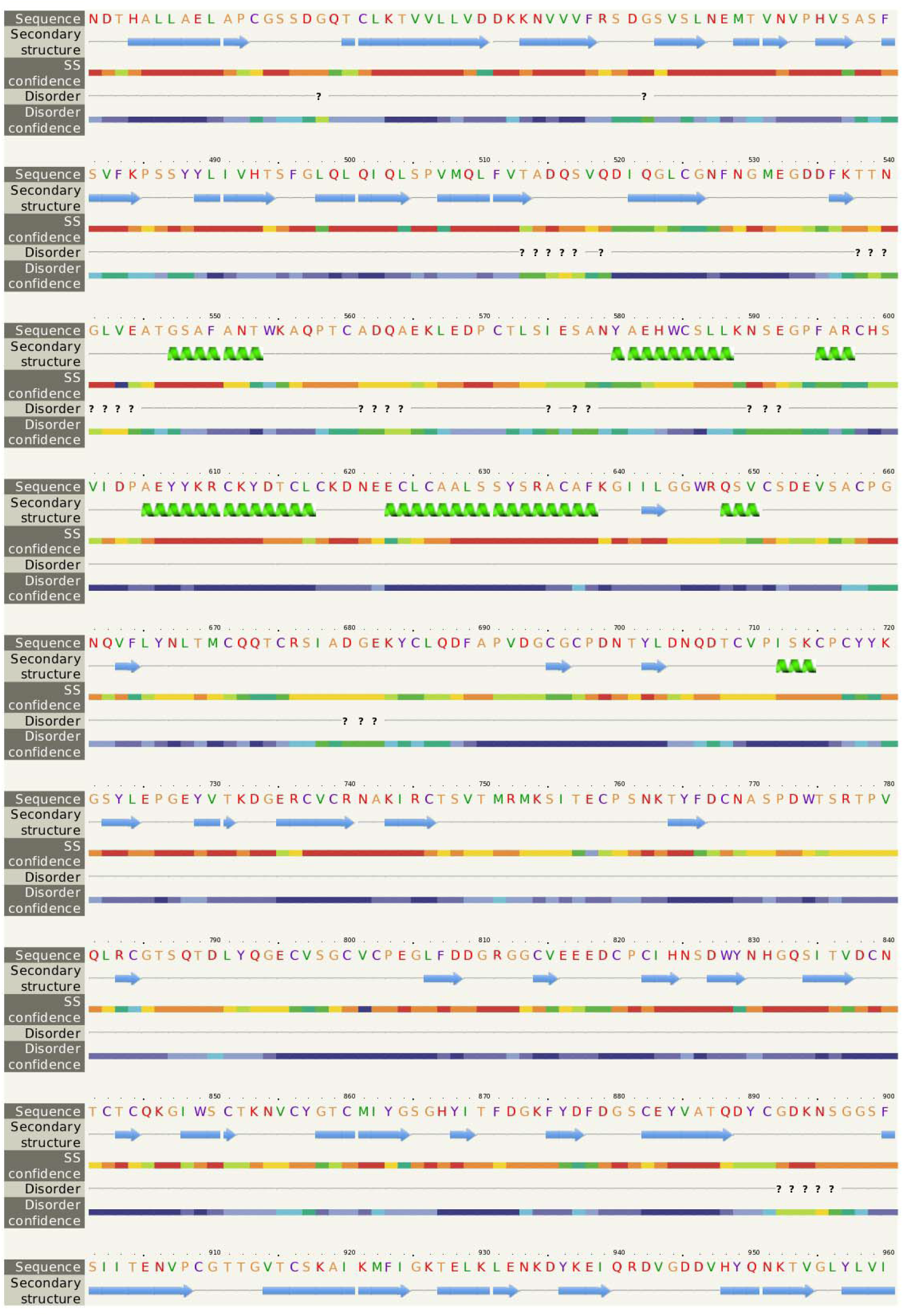
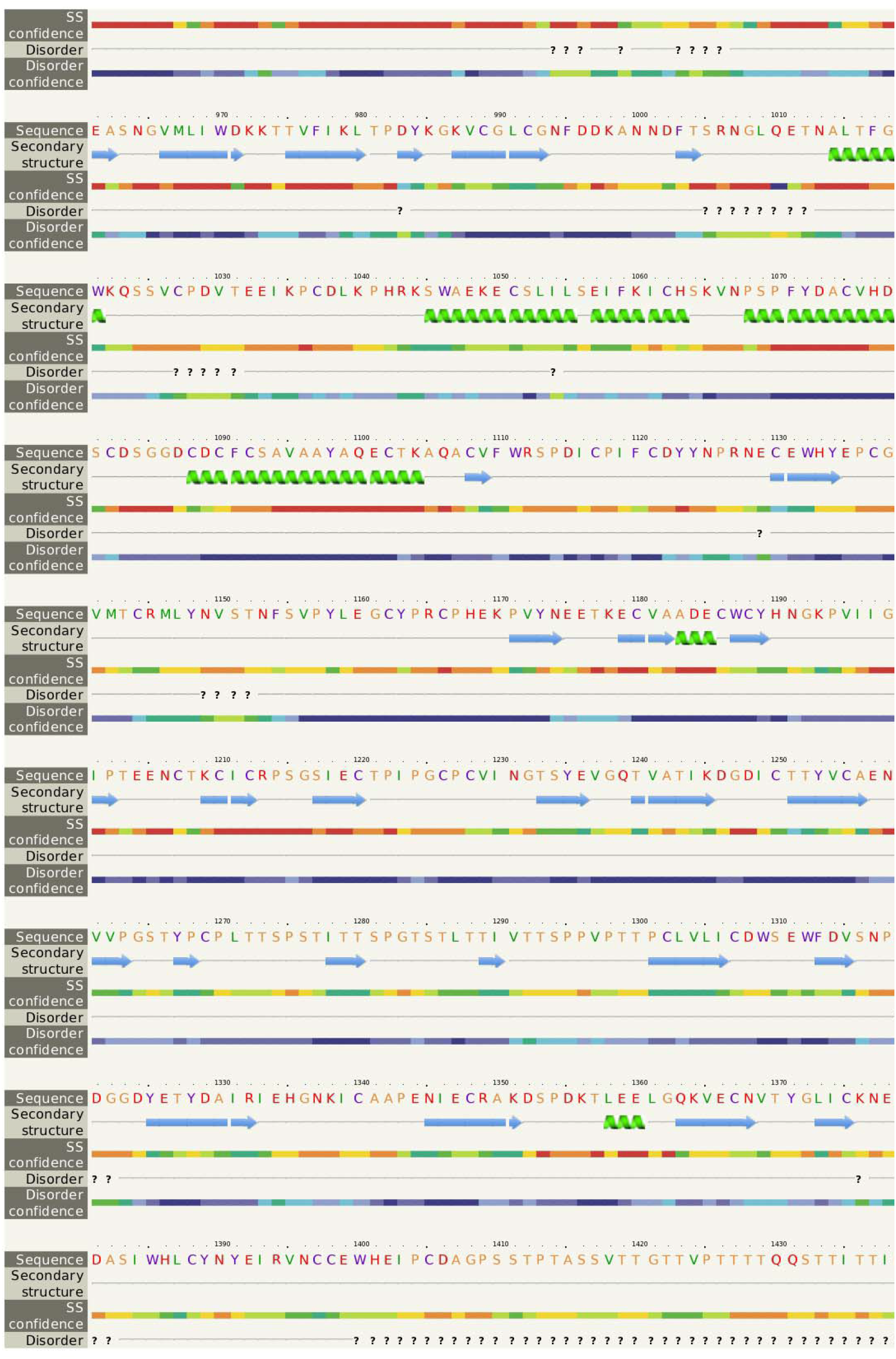
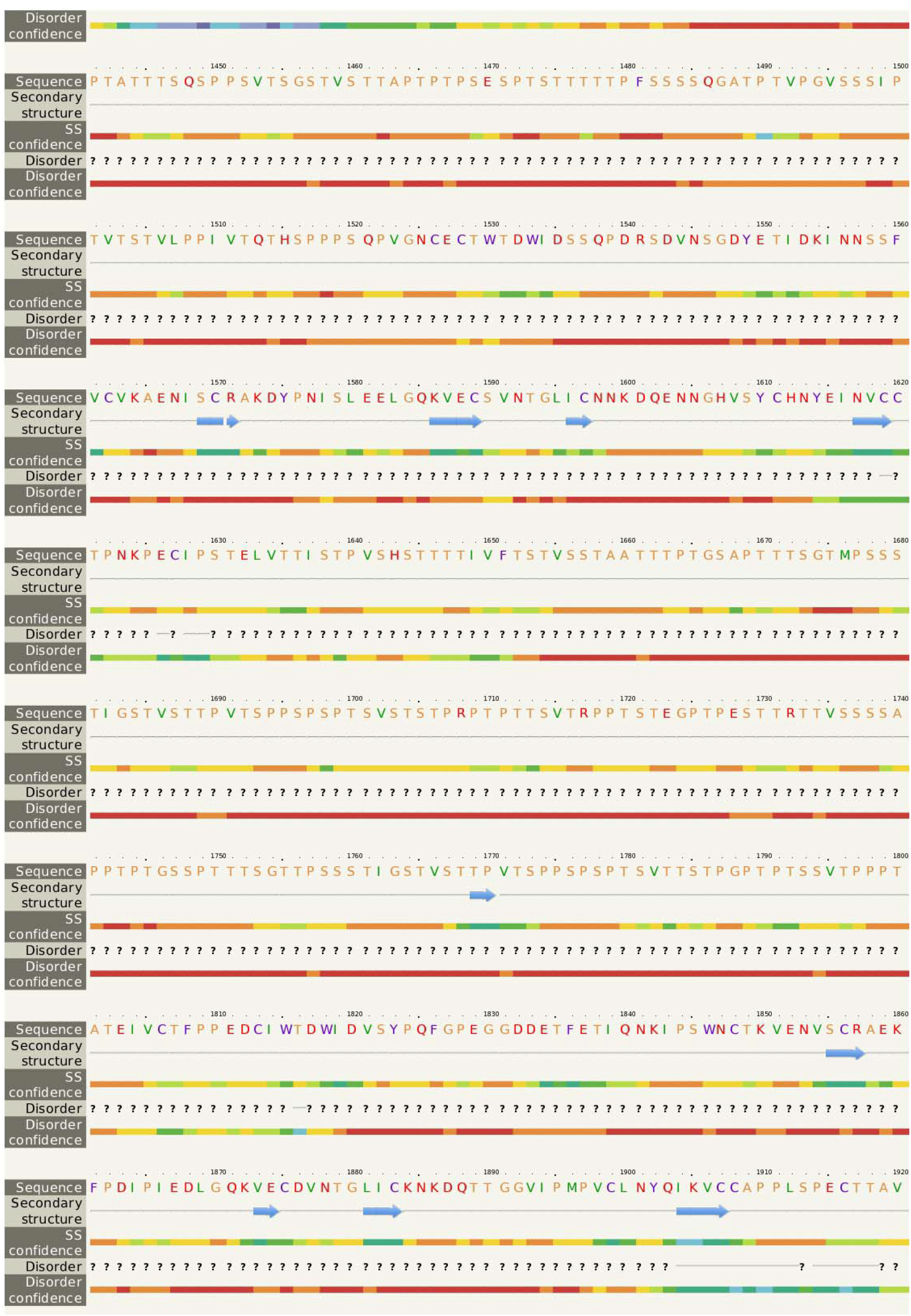
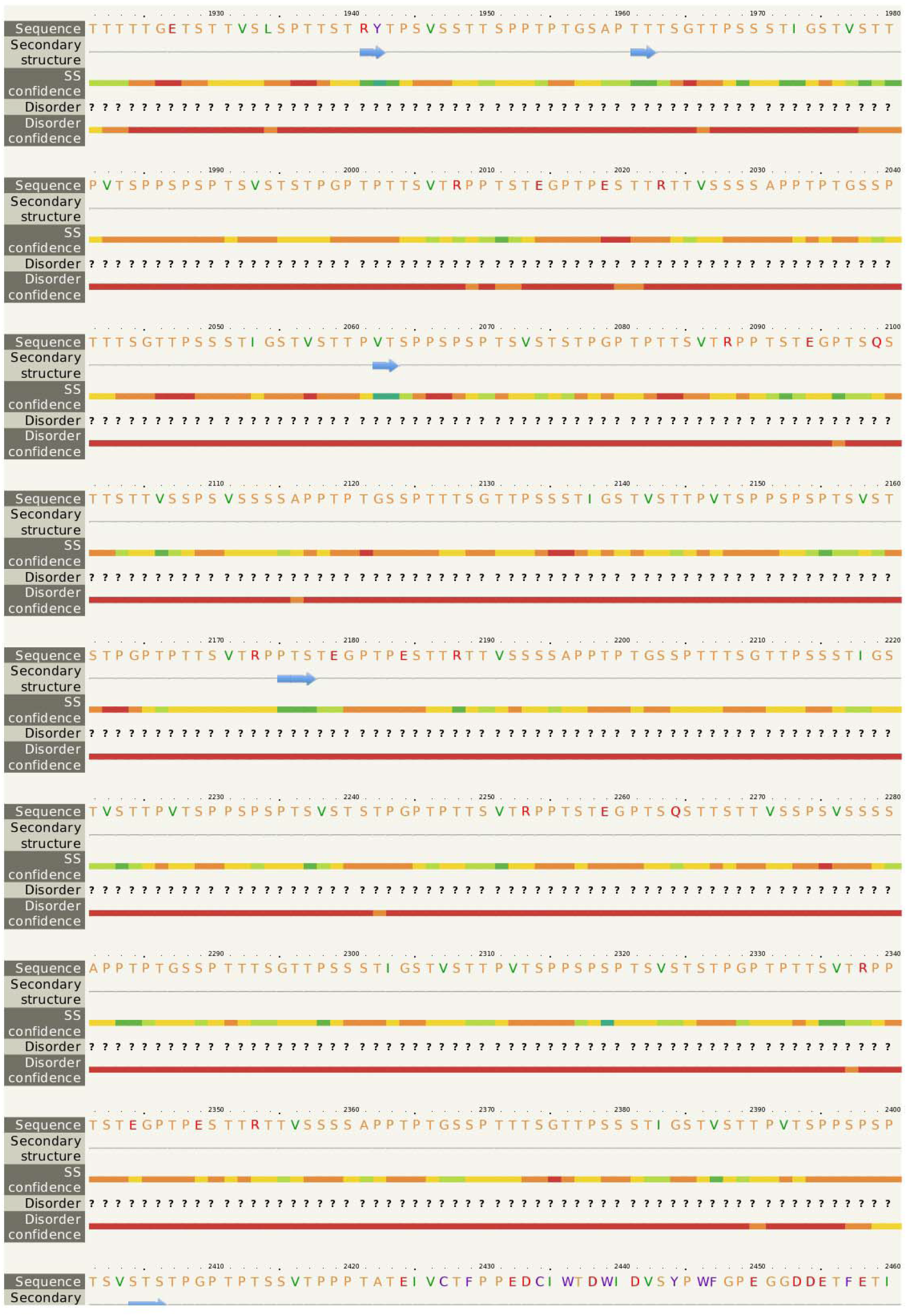
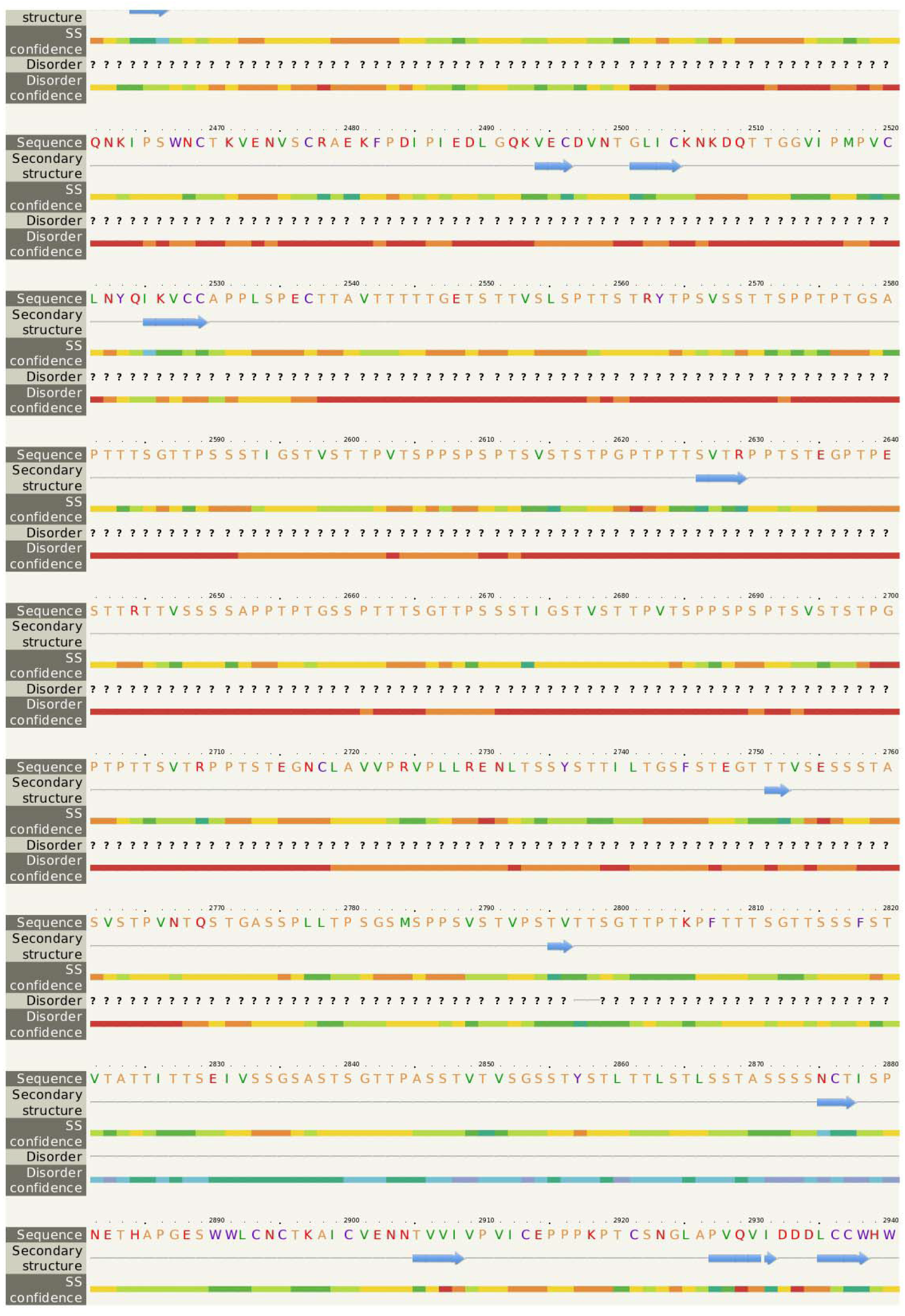
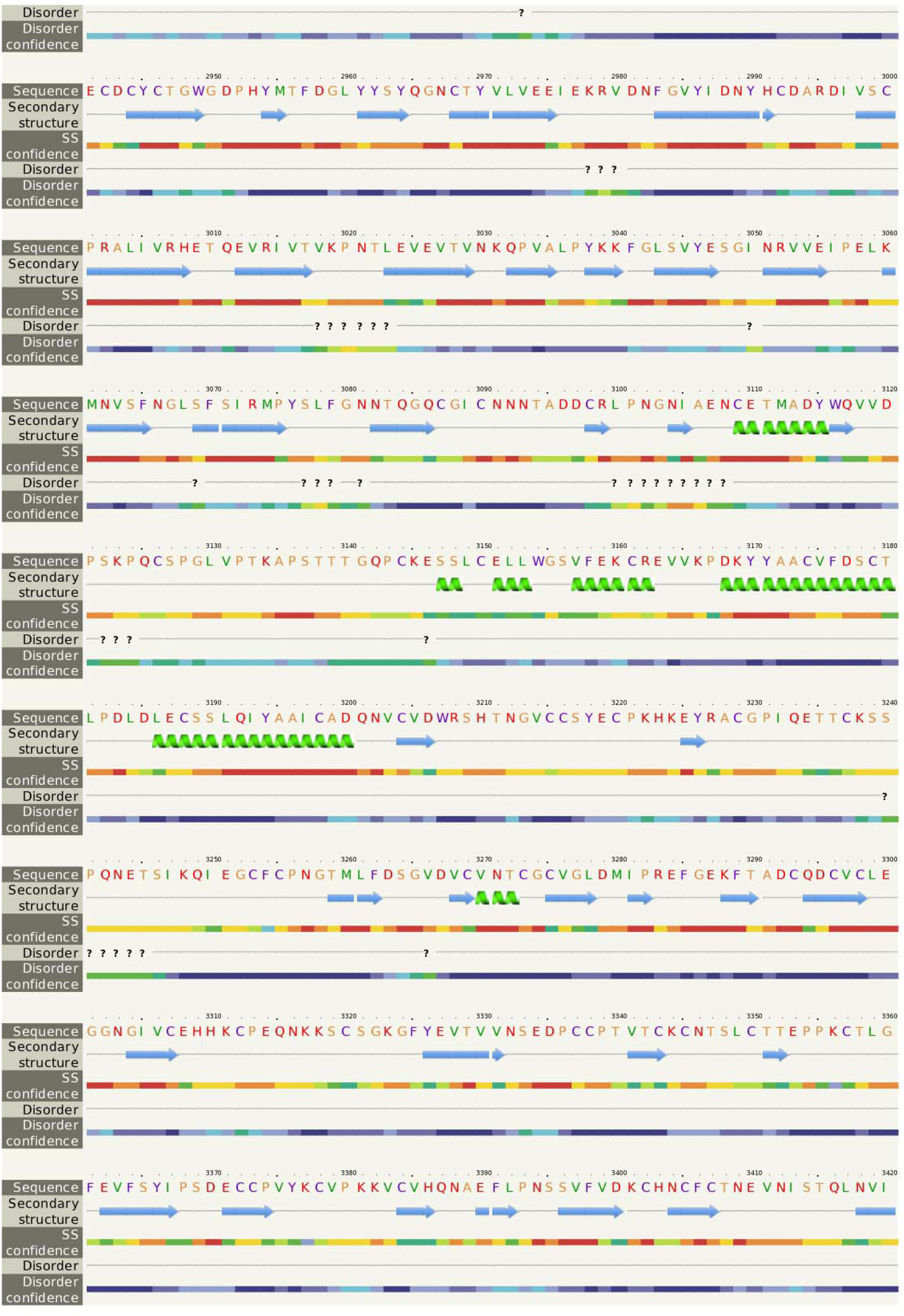
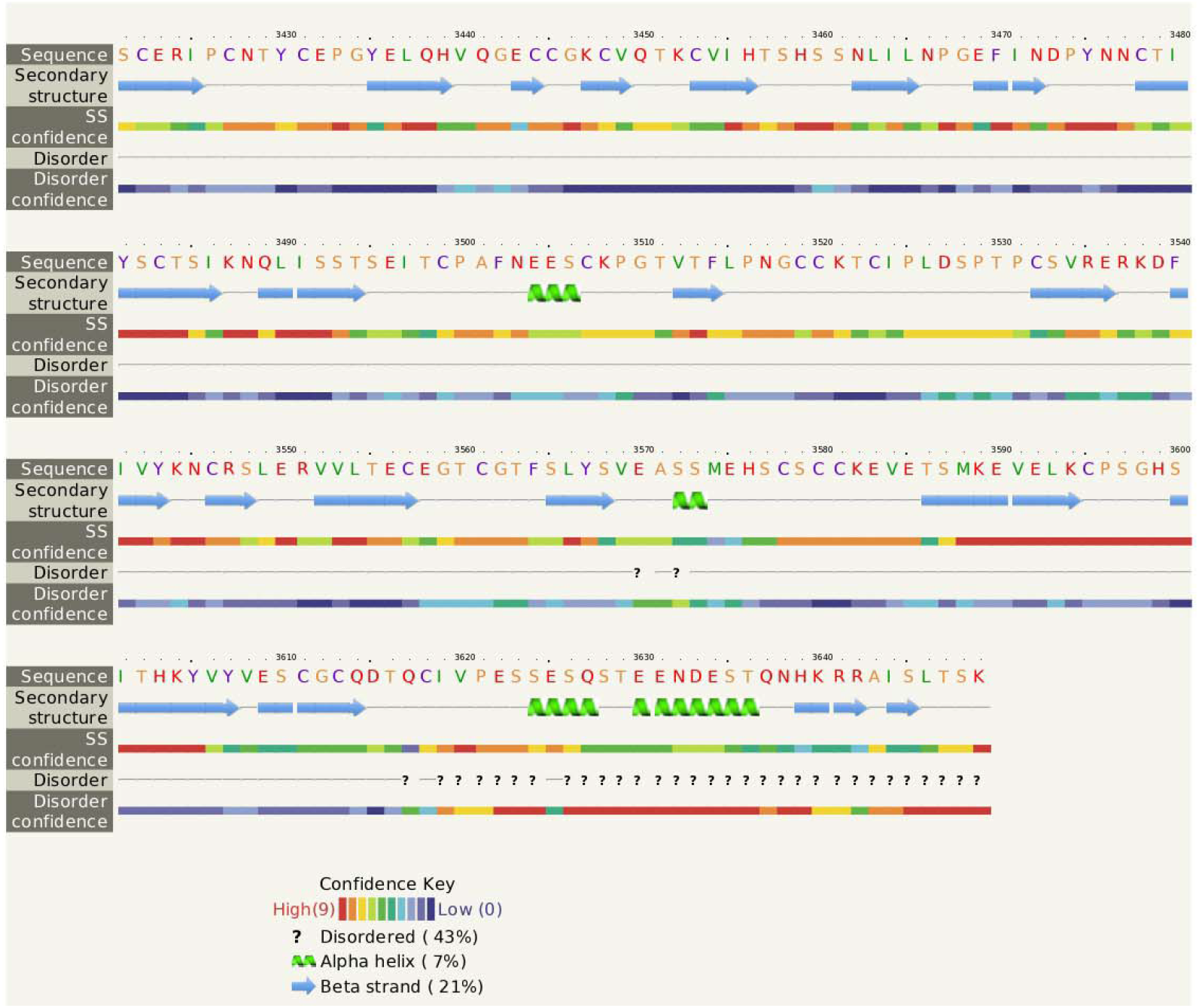
The pictorial description for secondary structure with the sites for alpha helix, beta sheet and loop structure for the mucin 2 gene of duck.

### Growth parameters for Bengal duck

In our current study, we had studied the growth parameters of indigenous duck of Bengal as Bengal duck. Body weight was studied of the ducklings just after hatching upto 10^th^ week of age (Table 1). **Fig 4** represents the diagrammatic representation of body weight at successive ages of brooding of Bengal duck upto 10^th^ week of age. Accordingly, average daily body weight gain was also recorded at successive ages up to 10^th^ week (Table 2). **Fig 5** shows the diagrammatic representation of the average daily body weight gain for both the groups of ducklings –Group I (Higher growth) and Group II (Lower growth). **Fig 6** represents the biomorphometric characteristics for adult Bengal duck from Group I (higher growth) and group II (Lower growth).

**Fig 4:**
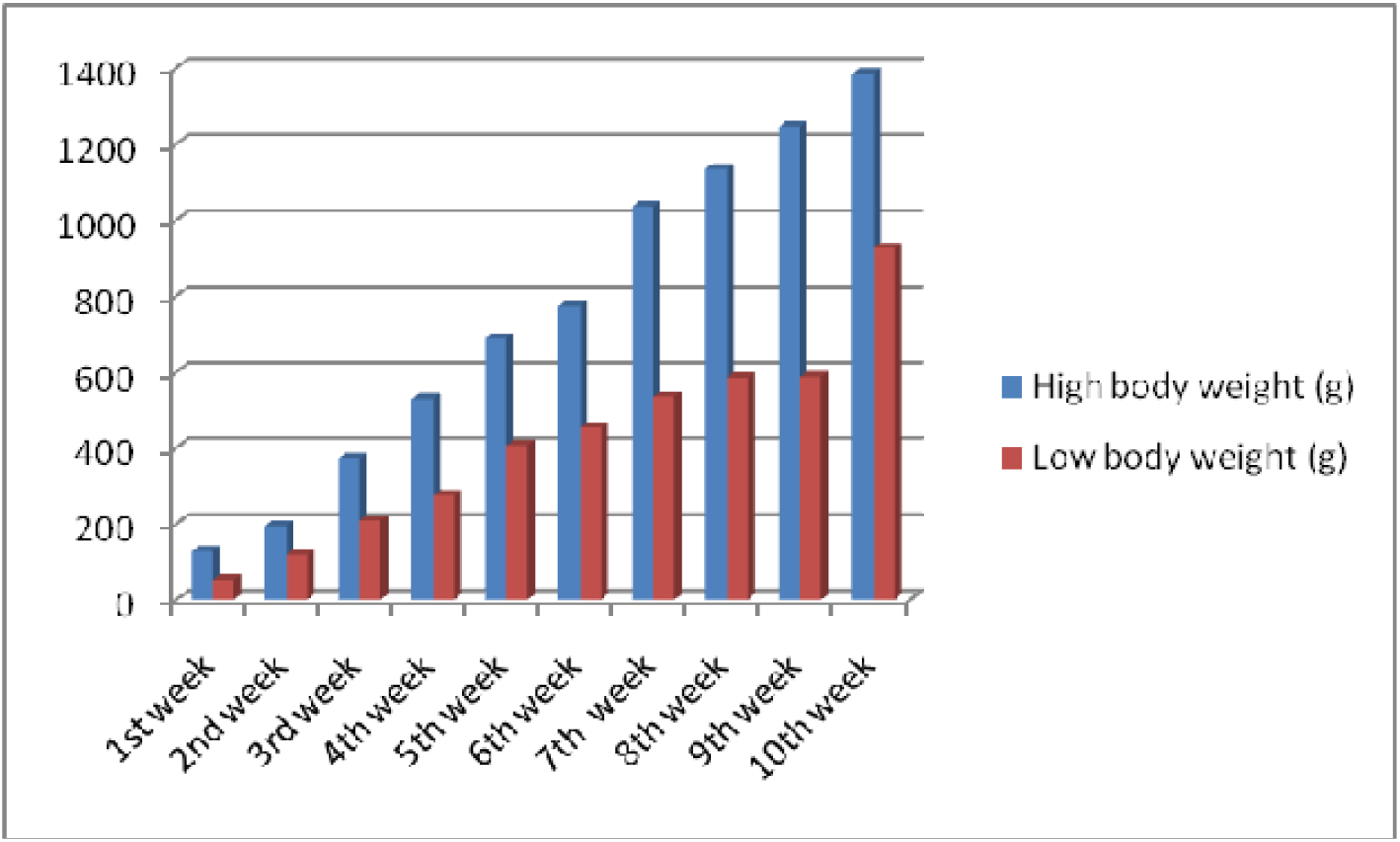
Graphical representation for body weight of indigenous ducks at successive stages of life

**Fig 5:**
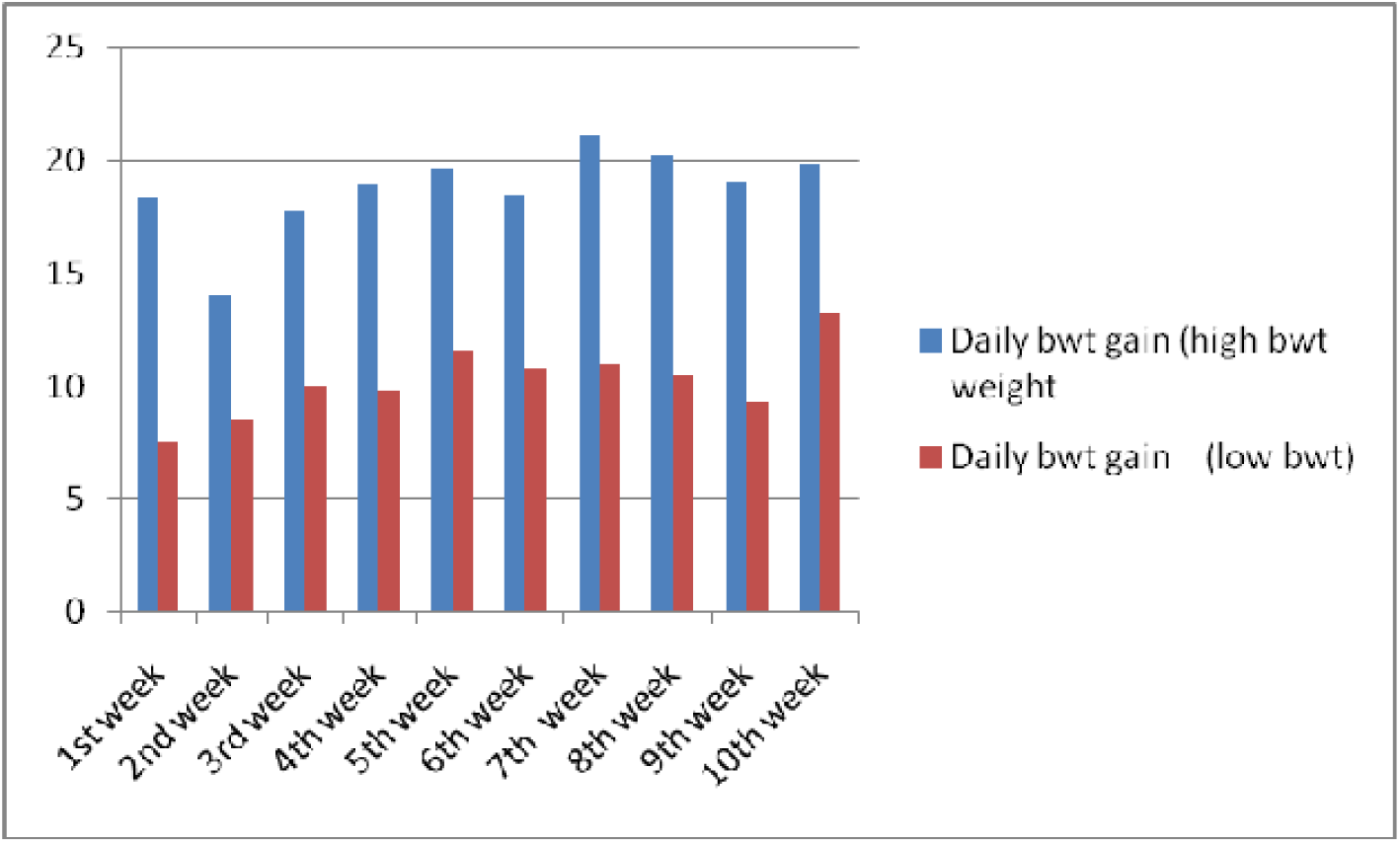
Daily body weight gain for Bengal duckling at successive ages

**Fig 6:**
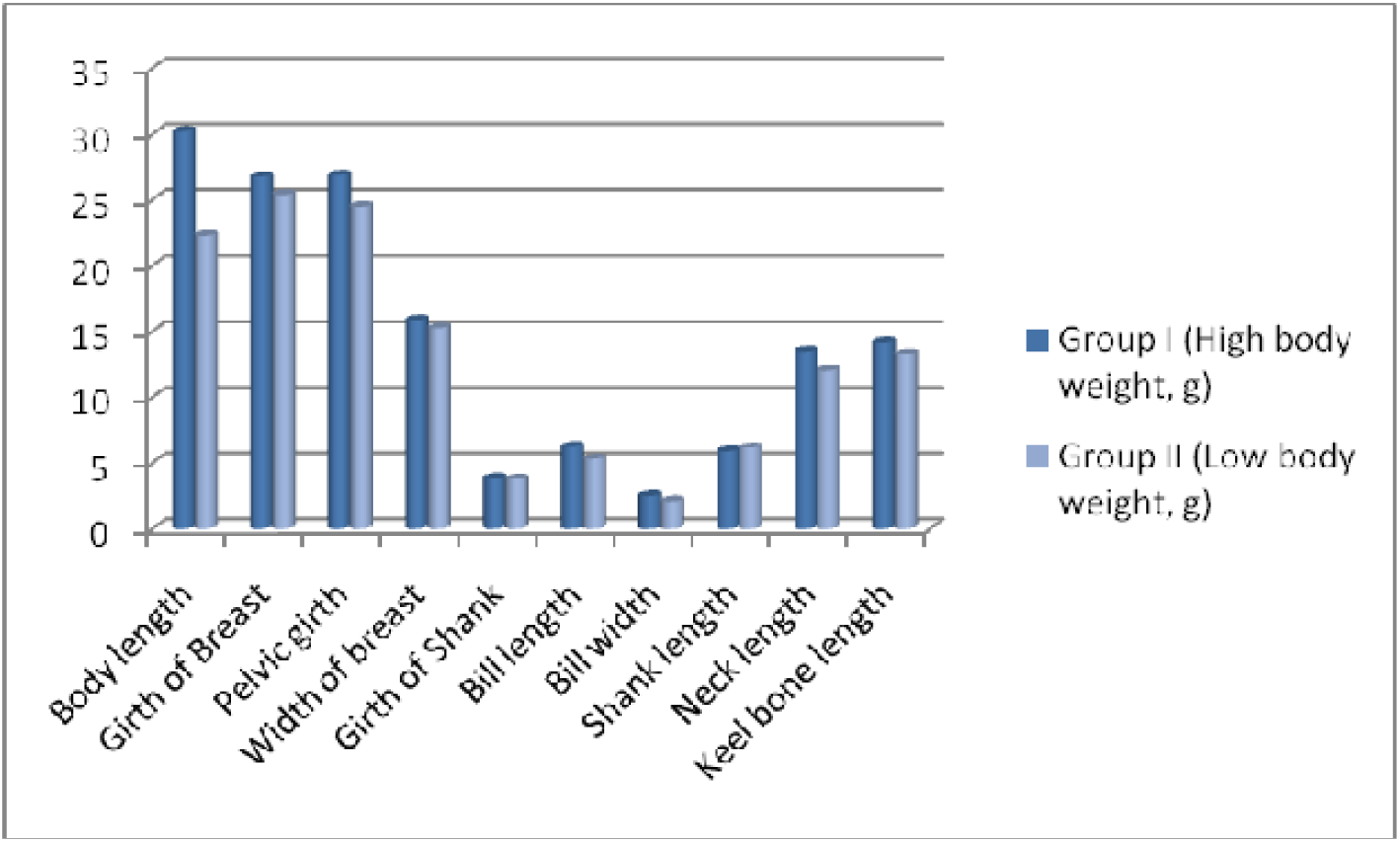
Biomorphometric characteristics for adult Bengal duck from Group I (higher growth) and group II (Lower growth)

**Table 2:**
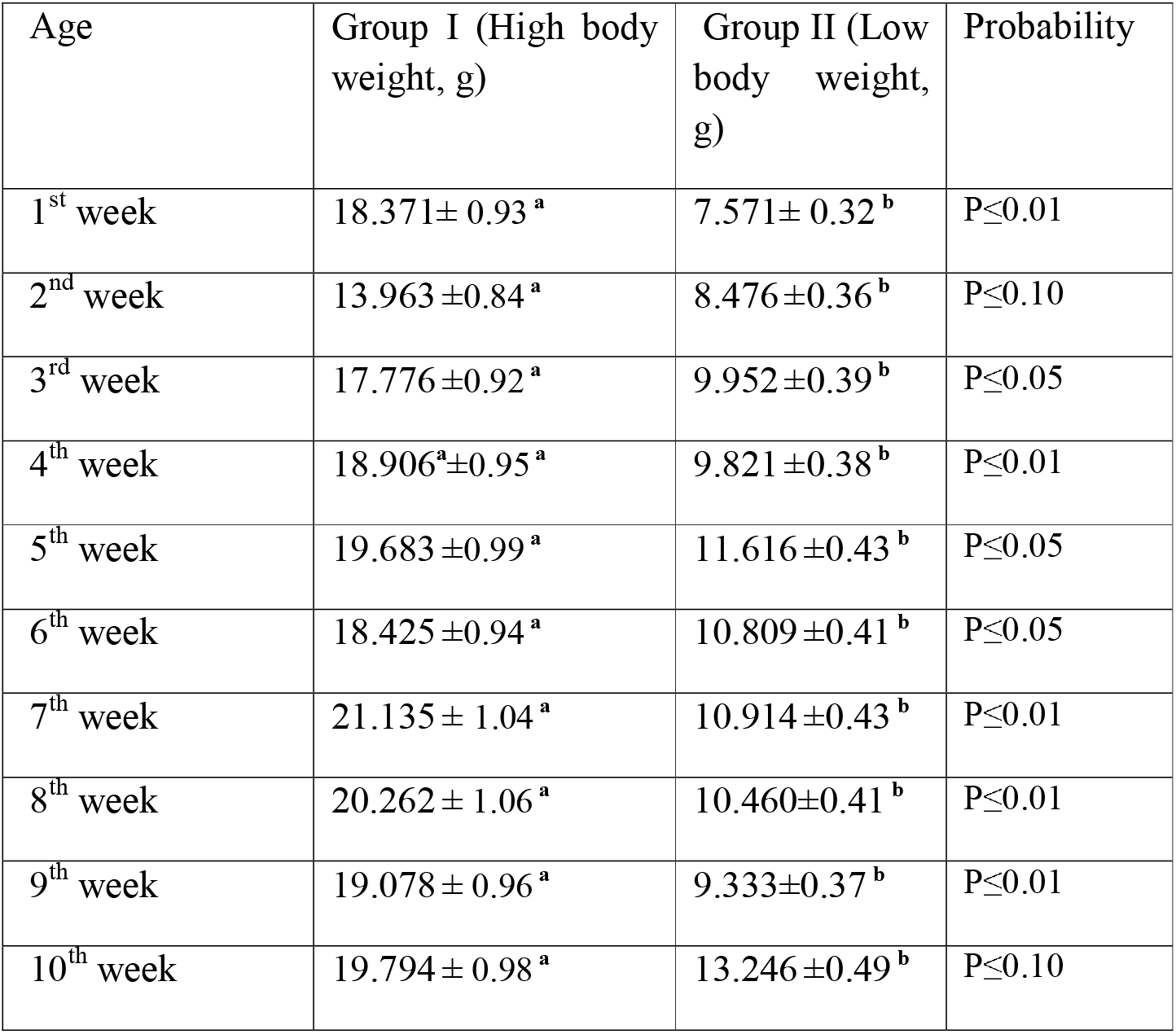
Average daily body weight gain.

| Age | Group I (High body weight, g) | Group II (Low body weight, g) | Probability |
| --- | --- | --- | --- |
| 1<sup>st</sup> week | 18.371± 0.93 <sup>a</sup> | 7.571± 0.32 <sup>b</sup> | $P \leq 0.01$ |
| 2<sup>nd</sup> week | 13.963 ±0.84 <sup>a</sup> | 8.476 ±0.36 <sup>b</sup> | $P \leq 0.10$ |
| 3<sup>rd</sup> week | 17.776 ±0.92 <sup>a</sup> | 9.952 ±0.39 <sup>b</sup> | $P \leq 0.05$ |
| 4<sup>th</sup> week | 18.906<sup>a</sup>±0.95 <sup>a</sup> | 9.821 ±0.38 <sup>b</sup> | $P \leq 0.01$ |
| 5<sup>th</sup> week | 19.683 ±0.99 <sup>a</sup> | 11.616 ±0.43 <sup>b</sup> | $P \leq 0.05$ |
| 6<sup>th</sup> week | 18.425 ±0.94 <sup>a</sup> | 10.809 ±0.41 <sup>b</sup> | $P \leq 0.05$ |
| 7<sup>th</sup> week | 21.135 ± 1.04 <sup>a</sup> | 10.914 ±0.43 <sup>b</sup> | $P \leq 0.01$ |
| 8<sup>th</sup> week | 20.262 ± 1.06 <sup>a</sup> | 10.460±0.41 <sup>b</sup> | $P \leq 0.01$ |
| 9<sup>th</sup> week | 19.078 ± 0.96 <sup>a</sup> | 9.333±0.37 <sup>b</sup> | $P \leq 0.01$ |
| 10<sup>th</sup> week | 19.794 ± 0.98 <sup>a</sup> | 13.246 ±0.49 <sup>b</sup> | $P \leq 0.10$ |

### Biomorphometric estimation for duck grouped into higher and lower growth parameters

Biomorphometric characteristics are useful parameters for assesing the growth parameters. They also serve as an indirect way to assess the body weight of the birds. The average biometry characteristics for adult male and female ducks including pool average for both higher (GroupI) and lower group (Group II) have been listed in Table 3. Significant differences in growth parameters were observed for Body length of Bengal duck. **Fig 6** represents the biomorphometric characteristics for adult Bengal duck from Group I (higher growth) and group II (Lower growth).

**Table 3.**
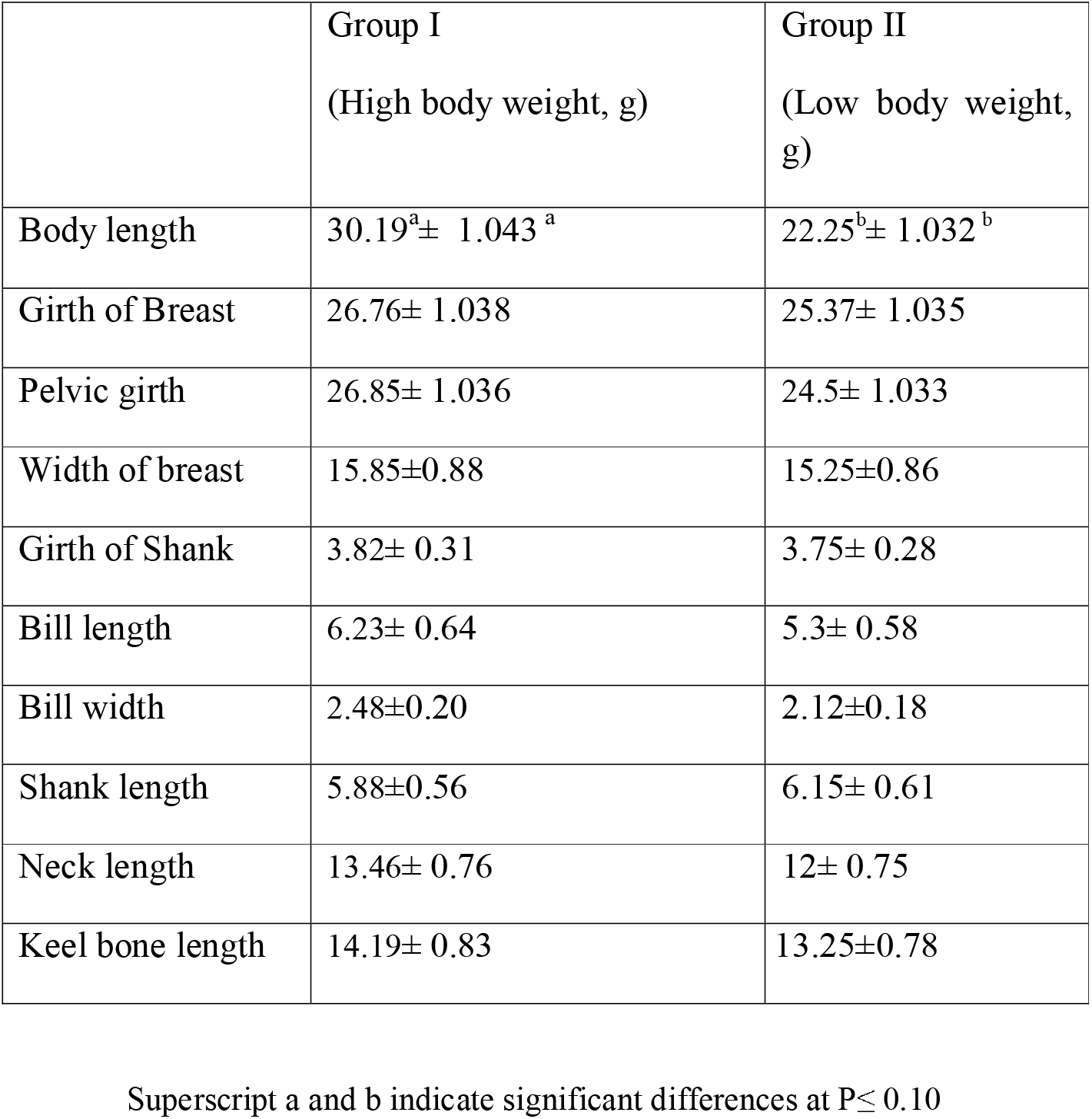
Biomorphometric characteristics for Bengal duck.

Growth curve for duck at successive stages of life based on Body weight is being represented at Fig 7, whereas Fig 8 depicts the growth trend with respect to daily body weight gain.

**Fig 7:**
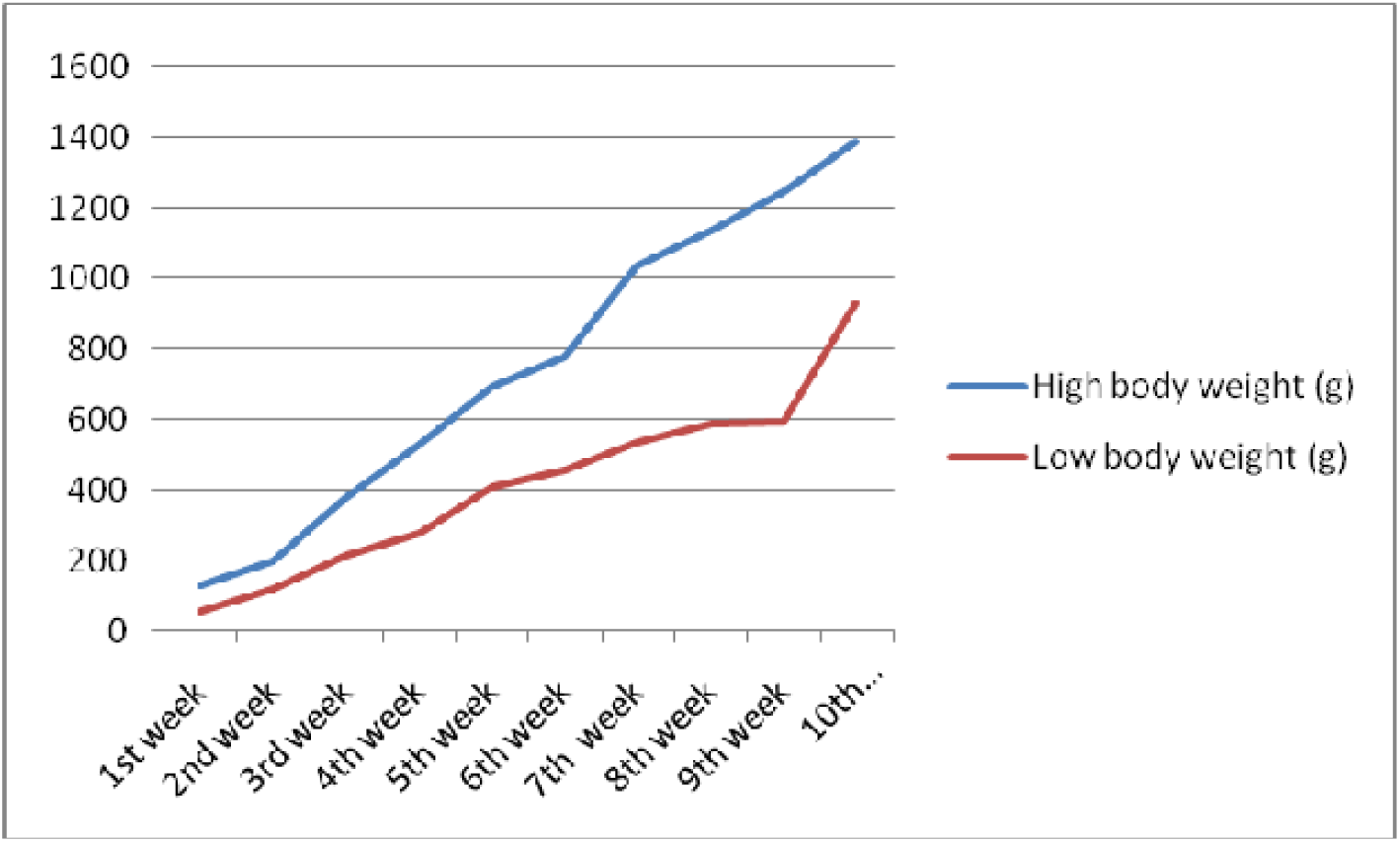
Growth curve for duck at successive stages of life based on Body weight.

**Fig 8:**
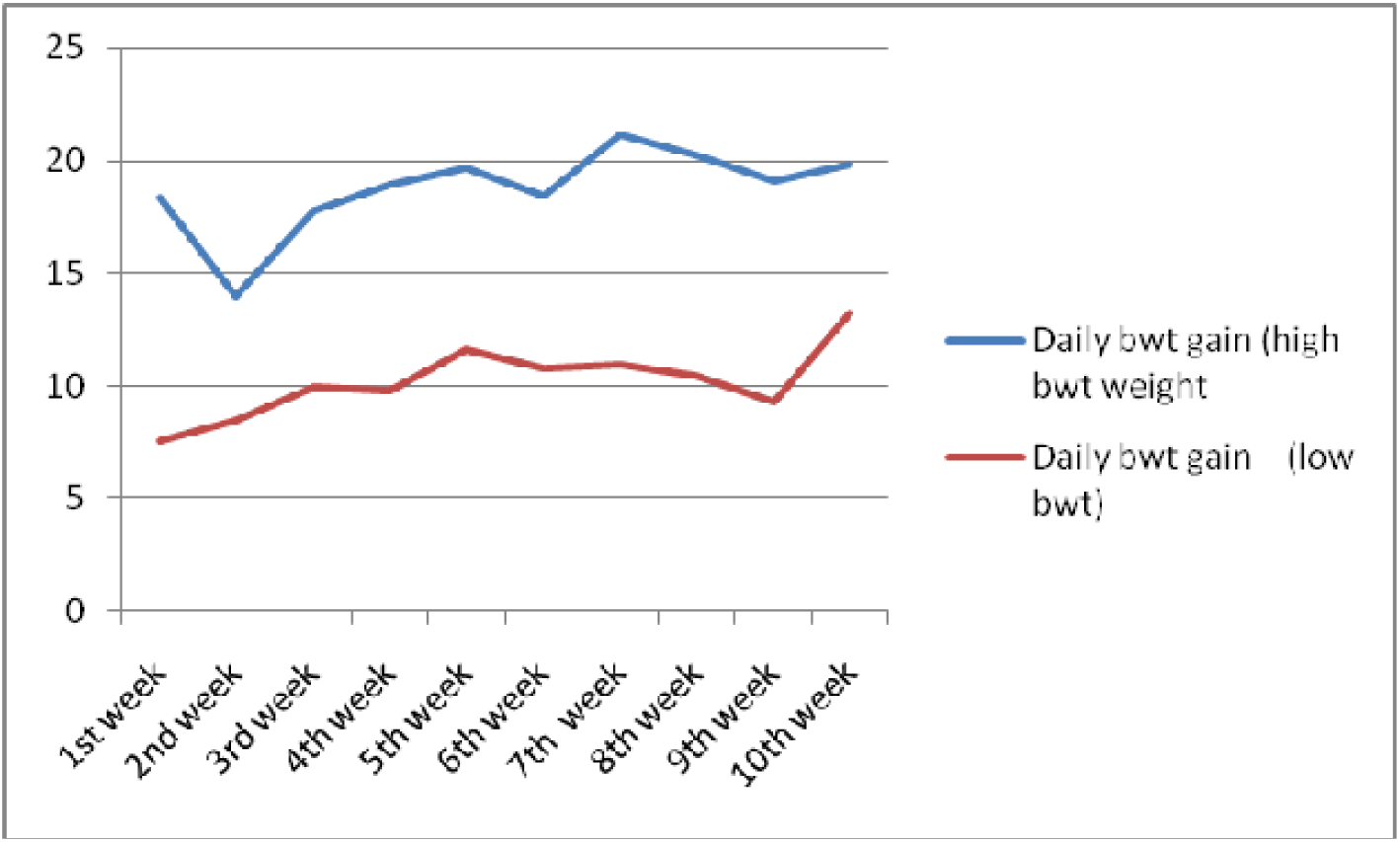
Growth trend representing daily body weight gain for Bengal duckling

### Expression profiling for Mucin 2 gene with respect to Growth parameters

Mucin is a protein expressed in the inner layer of epithelium of some organs. It is commonly observed in gut of livestock and poultry. In our current study, we studied the expression profiling of two important organs-duodenum (part of small intestinal) and caecum (part of large intestinal) of Bengal duck with respect to high and low body weight.

Fig 9 and Fig 10 reflects the better expression profile for mucin gene in Group I compared to that of Group II for duodeum and caecum respectively. The expression for mucin gene was observed to be more than five folds in Group I compared to that of Group II. It is evident from Fig 9 that although a similar trend of better expression of mucin is observed for both the organs for gut, mucin is better expressed in duodenum compared to that of caecum. The line graph represents steep line in mucin expression in duodenum, compared to that of caecum. The expression of mucin gene in Group I was observed to be more than six folds (6.29) in duodenum compared to that of caecum. Similarly, expression of mucin gene was observed to be more than five folds (5.3) in duodenum compared to that of caecum. Fig 11 represents the consolidated expression profiling for duck mucin2 gene in the gut of duck.

**Fig 9:**
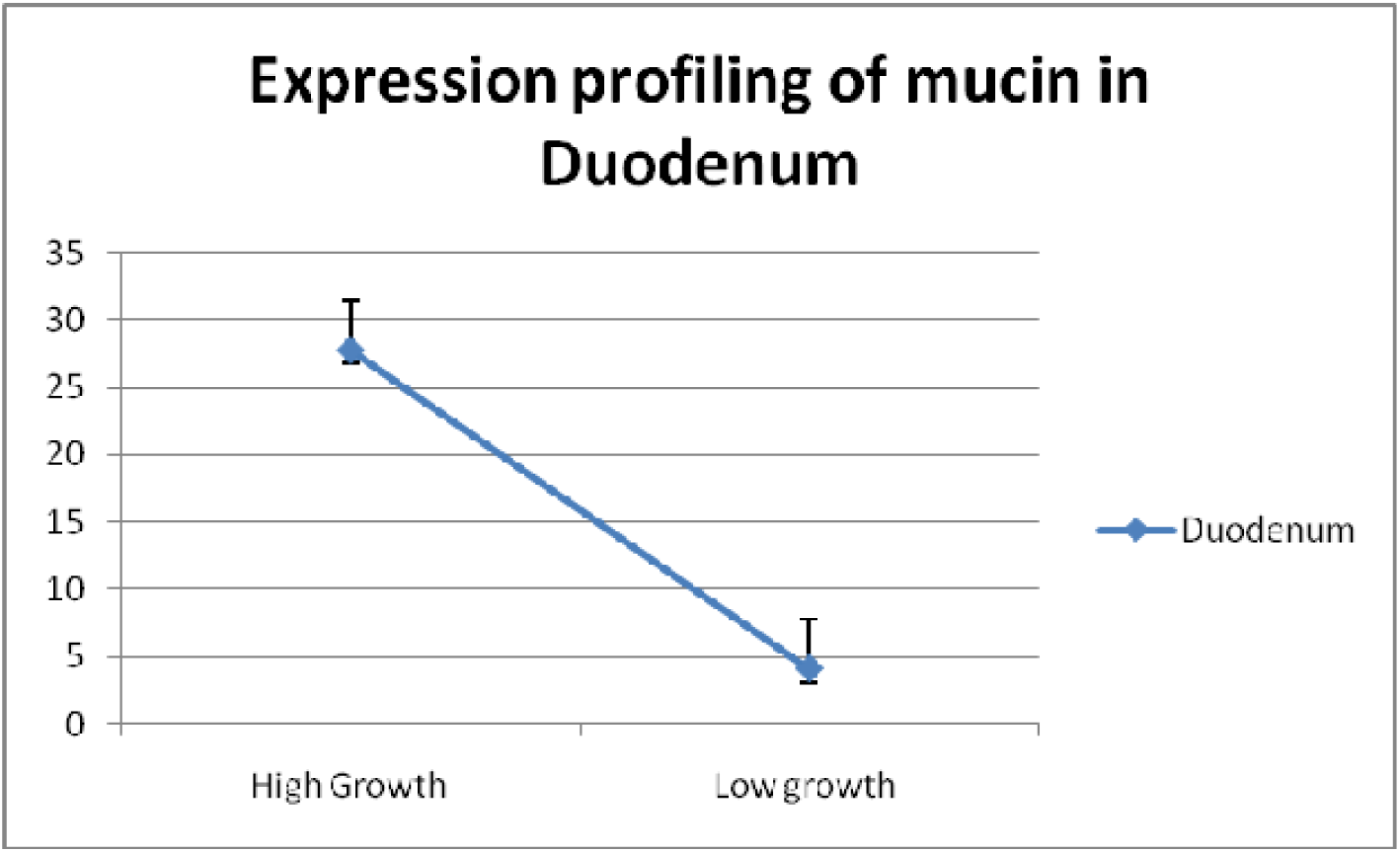
Expression profiling for Mucin 2 gene with respect to Growth parameters in duodenum

**Fig 10:**
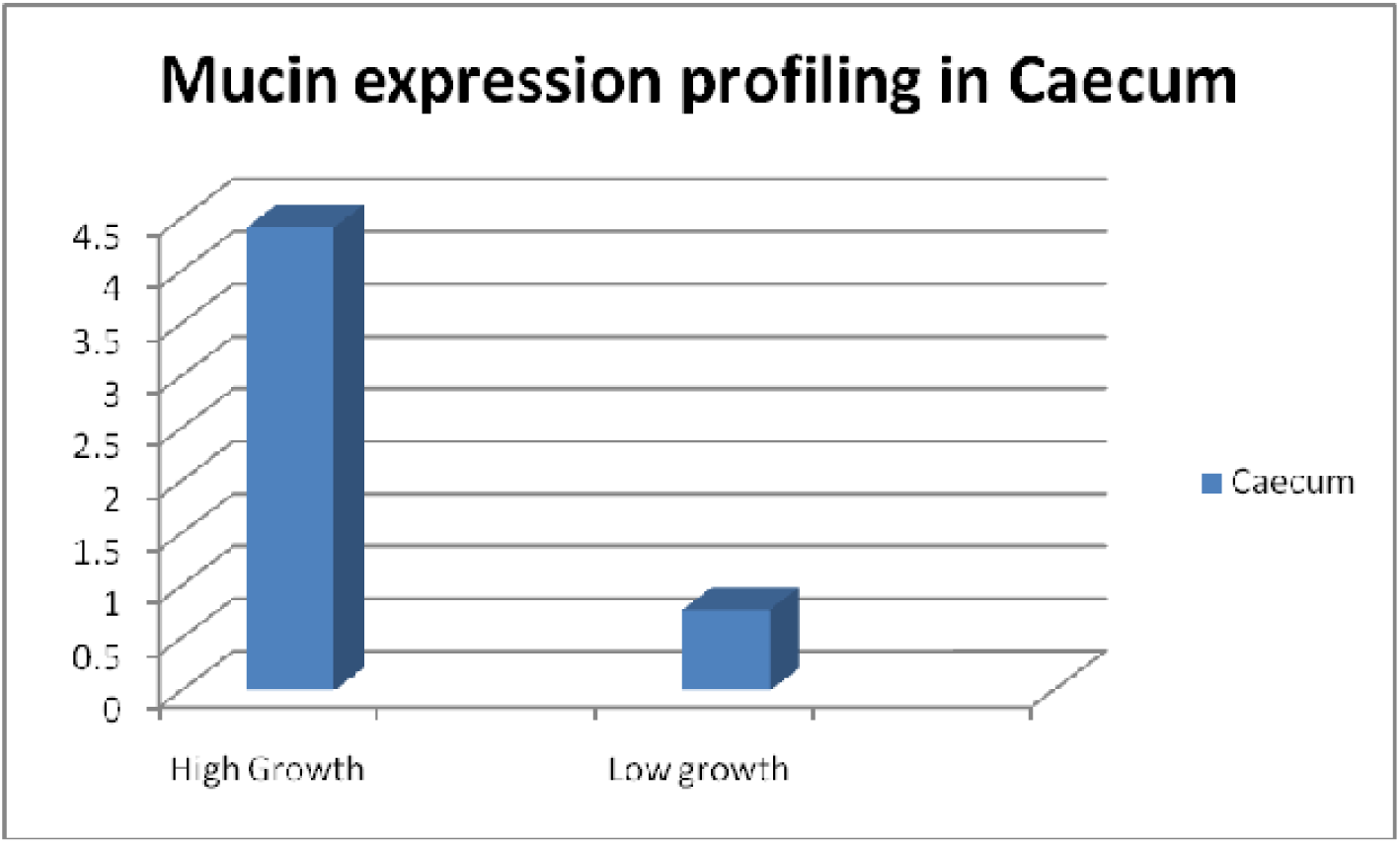
Expression profiling for Mucin 2 gene with respect to Growth parameters in caecum

**Fig 11:**
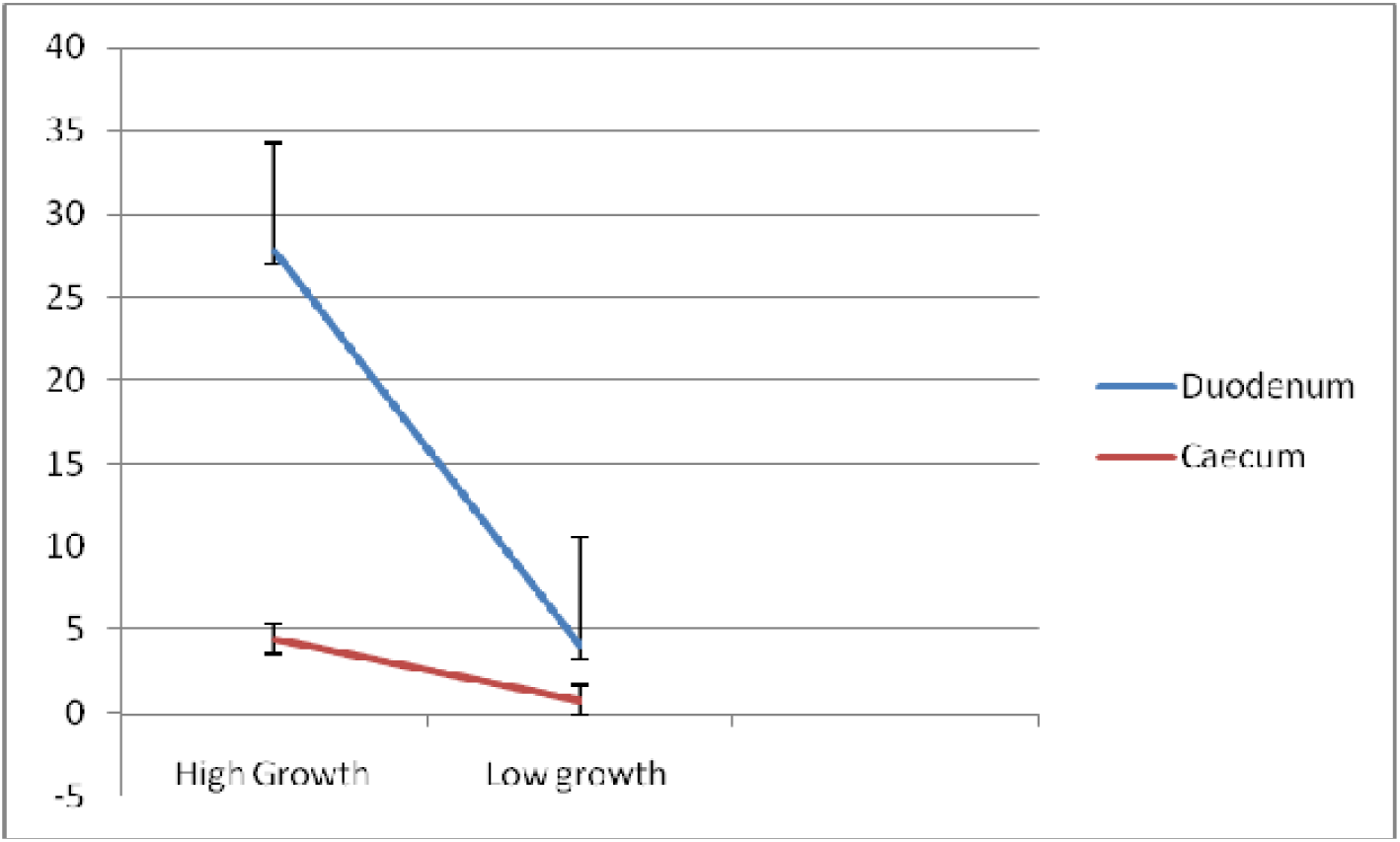
Differential mRNA expression profiling for Mucin2 gene in gut of Bengal duck

## Discussion

Mucus is a viscous gel-like material covering the gastro-intestinal mucosal surface. The entire surface of the chicken gastrointestinal tract is covered by a layer of mucus that functions as a diffusive barrier between the intestinal lumen and absorptive cells. The mucins are the main component of the mucus layer, which produces and secreted by goblet cells. Mucin from duck is sequenced and characerized for the first time, hence comparison was not possible. We observed certain important domains of duck mucin as VWFD (VonWillebrand factor) domain, VWFC-1, VWFC-2 domains, CTCK1 (C terminal cystine knot signature), CTCK2 (C terminal cysteine knot domain profile)

The structures of transforming growth factor-β (TGF-β), nerve growthfactor (NGF), platelet-derived growth factor (PDGF) and gonadotropin have beenshreferences therein]: these proteins are foldedinto two highly twisted antiparallel pairs of β-strands and contain threedisulfide bonds, of which two form a cthird bondpasses (see the schematic representation below). This structure is calledcystine knot to the cystine-knot family [3] and which istherefore called C-terminal cystine-knot (CTCK). Members of the C-terminalcystine knot family are listed as-von Willebrand factor (vWF), a multifunctional protein which is involved in maintaining homeostasis. It consists of 4 vWF type D domains, 3 vWF tydomains, 2 vWF type C domains (see <PDOC00928>, aX domain and the C-terminal cystine knot.

Mucins. Human mucin 2, a highly polymorphic multidomain molecule with a modular architecture similar to vWF. Xenopus mucin B.1 which contaiC domain, a X domain and a CTCK. Other mucins thatcontain a CTCK are the human tracheobronchial mucin (gene MUC5), bovinesubmaxillaryapomucin and rat intestinal mucin-likeprotein. CCN family (cef-10/cyr61/CTFG/fisp-12/nov protein family). These growth-factor inducible proteins are structurally related to the insulin-likegrowt<PDOC00194>) and could also function asgrowth-factor binding proteins. Drosophila slit protein which is essential for development of midline gliaand commissural axon pathways. It is composed of four leucine-rich repeaa laminin G-like repeat and the CTCK.

Norrie disease protein (NDP) which may be involved in neuroectodermal cell-cell interaction and in a pathway that regulates neural cell differentiation (https://prosite.expasy.org/doc/PS01185). Von Willebrand factor (VWF) is a large, multimeric blood glycoprotein synthesized in endothelial cellsmegakaryocytes, that is required for normal hemostasis. The type D domain (VWFD) is not only require clotting factor VIII binding but also for normal multimerization of VWF (https://prosite.expasy.org/PDOC00928). The VWFC domain is named after the von Willebrand factor (VWF) type C repeat which is found twice in this multidomain protein. It has a length of about 70 amino acids covering 10 well conserved cysteines (https://prosite.expasy.org/rule/PRU00580). von Willebrand factor for which the duplicated VWFC domain is thought to participate in oligomerization, but not in the initial dimerization step. The presence of this region in other complex-forming proteins leads to the assumption that the VWFC domain might be involved in forming larger protein complexes.

We could identify an important domain involved with growth as EGF1 (EGF like domain signature), presumed to play a major role in growth promotion. Certain important observations were revealed that since disulphide bonds were not included in this region, it is presumed to be open. EGF1 domain is inserted within VWFC-1 and VWFC-2. These are presumed to effect growth traits (Hunt and Barker, 1997, Bork., 1993, Bork., 1991, Voorberg et al., 1991). Mucins are high-molecular weight glycoproteins (50–80% O-linked oligosaccharides) produced by epithelial tissues in most animals. These glycoproteins are found in mucus (e.g. saliva, gastric juice, etc.) and secreted by mucous membranes to lubricate or protect body surfaces of vertebrates and they have a central role in maintaining epithelial homeostasis (Marín *et al*., 2012). Since this is the first report of studying mucin gene expression with respect to growth parameters in any livestock species, comparison as such is not possible. However, there exists certain reports which depicts that mucin is responsible for growth.

The mucus layer is part of the innate host response, protecting against luminal microflora, preventing gastrointestinal pathologies, and participating in the processes of nutrient digestion and absorption (Forstner *et al*., 1995). Decrease in mucin synthesis in poultry could compromise the mucus layer and reduce nutrient utilization (Horn *et al*., 2009). In addition to its protective functions, mucin is involved in filtering nutrients in the gastrointestinal tract (GIT) and can influence nutrient digestion and absorption (Montagne *et al*., 2004). Any component, dietary or environmental, that induces changes in mucin dynamics has the potential to affect viscosity, integrity of the mucus layer, and nutrient absorption. In the current study, we observed better expression of mucin 2 gene in duodenum in comparison to that of caecum. Since it is evident that duodenum is greatly involved with the function of absorption of nutrients, better expression reveals that mucin 2 may be involved in nutrient absorption.

Hence it is evident that mucin has an active role in the process of nutrient digestion and absorption & utilization. Thus, in the duck expression of less mucin indicate less nutrient content in the body which will ultimately affect the growth. Mucins in general contain many threonine and serine residues, which are extensively O-glycosylated. Due to this profound glycosylation, mucins have a filamentous conformation. Reports are also available indicating mucin aids in threonine absorption (Toribara *et al*. 1993). We observed high glycosylation of both O-linked and N-linked glyosylation.

Certain set of reports indicate the role of threonine in growth. Debnath *et al*. (2018) studied that threonine supplementation can positively influence antioxidant enzyme activities and haemato-biochemical parameters in commercial layers. Because threonine is one of the essential amino acids liable to be limited when high humidity heat stress would decrease feed intake. Although no direct report is available about the mechanism of mucin in growth.

Certain reports are available for differential expression profiling of mucin gene in other aspects. It has been reported that probiotic supplementation increased the expression of the MUC2 gene in the chicken jejunum (Smirnov et al., 2005) and rat colon (Caballero-Franco et al., 2006). Different factors such as microbial colonization in the intestine can affect the production, secretion and composition of mucin (Forder *et al*., 2007; Azzam *et al*.,2011). Aliakbarpour *et al*., (2012) studied that Inclusion of lactic acid bacteria-based probiotic in the diets significantly increased goblet cell number and villus length (p<0.05) and significantly increased gene expression (p<0.05) with higher intestinal MUC2 mRNA in birds fed diet with probiotics.

Mucin is a protein expressed in the inner layer of epithelium of some organs. It is commonly observed in gut of livestock and poultry. In our current study, we studied the expression profiling of two important organs-duodenum (part of small intestinal) and caecum (part of large intestinal) of Bengal duck with respect to high and low body weight.Better expression profile for mucin gene in Group I compared to that of Group II for duodeum and caecum respectively was observed. The expression for mucin gene was observed to be more than five folds in Group I compared to that of Group II. Although a similar trend of better expression of mucin is observed for both the organs for gut, mucin is better expressed in duodenum compared to that of caecum. The line graph represents steep line in mucin expression in duodenum, compared to that of caecum. The expression of mucin gene in Group I was observed to be more than six folds (6.29) in duodenum compared to that of caecum. Similarly, expression of mucin gene was observed to be more than five folds (5.3) in duodenum compared to that of caecum.

Since this is the first report of studying mucin gene expression with respect to growth parameters in any livestock species, comparison as such is not possible. However, there exists certain reports which depicts that mucin is responsible for growth. We observed significant differences of body weight and body weight gain at successive stages of age. We similarly observed significant differences in body length of the Bengal duck among the biomorphometic traits under consideration. Lack of significant differences in other biomorphometric traits may be due to the inherent breed chracteristics of Bengal duck. As explained earlier that we had classified the entire duckling population data into high body weight (better growing) as values above mean +standard deviation (***x*** □ **+SD**) and low body weight (less growing) as values less than mean –standard deviation (***x*** □ **−SD**). Growth curve for duck at successive stages of life based on Body weight is being represented in Fig 7, whereas the trend for average daily body weight gain have been depicted in Fig 8.

It has been observed that the body weight of the ducks at brooding stage increases at significantly from lower body weight group of ducklings. However, the highest difference was observed at first week of age. An increase of 58.78 percentage was observed in higher group in comparison to lower bwt. group. In the duckling of lower body weight group, from 8^th^ week to 9^th^ week, body weight was recorded to be similar. This is reflected as decrease in daily body weight gain at 9^th^ week of age. It is interesting to note that similar trend was observed for the ducklings of higher body weight group. Average daily body weight was observed to be lowered at 9^th^ week of age. Since this is the first report of body weight estimation and rearing of duckling of Bengal duck (indigenous duck of West Bengal), comparison was not possible as such. We predict certain physiological changes in the duck at this stage of life, as the body diverts a part of nutrients for the development of reproductive organs.

Average daily body weight gain was observed to be decreasing at second week of age at higher body weight group. An increasing trend was observed upto 5^th^ week of age. To compare the growth performance of Bengal breeds of ducks with other available duck genetic resources average weekly live weight of Nageswari, Pekin, Muscovy and Desi white ducks were collected from a published paper of Morduzzaman *et al*. (2015) and Bhuiyan *et al*. (2005). Body weight for Nageswari ducks were reported to be slightly more than Bengal duck as for ducklings from sucessive week of age. This variation might be due to the differences in feed availability, nutrients content in feed, management practices and selection of the ducklings. Among these breeds including desi duck of West Bengal, the highest body weight was produced by Pekin breeds of ducks (1763 gm) and lowest in Desi white (1208 gm). Gonzalez and Marta (1980) observed that the body weight of female khaki Campbell ducklings at 1, 4 and 7 weeks of age averaged 85.6, 585.1 and 1113 gm, while that of males were 75.0, 594.4 and 1213.3 gm for the same period. Body weight of White Pekin ducklings at 6^th^ week of age averaged 1350 gm and at 7^th^ week 1718 gm (Aggarwal *et al*., 1981). In another investigation Andrews *et al*., (1984) reported adult body weight of desi ducks under intensive and semi-intensive system of rearing was 1311 and 1281 gm respectively.

The data on the mean daily body weight gain of indigenous ducks indicated that the body weight was increased highest at 7^th^ week and after 7^th^ week the body weight is decreased little by little. The body weight at all ages were significantly (p<0.05) different between generations. Higher mean value (11.4g/bird) for daily body weight gain was recorded for those birds which fed wet mash than the birds which fed dry mash Ogbonna *et al*. (2000). Bale-Therik and C. Sabuna (2010) reported that the bird fed with grit, increase the feed intake, it stimulated the digested enzyme and improve the body weight. There was an average daily growth rate of 5.88 ± 0.05 and 6.27 ± 0.09 g per bird per day at their 8th weeks growth phases, respectively (Faruque *et al*. 2013).

But certain significant variations were detected in the phenotypic data for growth at different ages of duck. In the next step, we attempt to explore that if any quantitative variations are present in mucin gene. These quantitative expression profiling may arise due to promoter region of the gene.

### Conclusion

It can be concluded that genetic variability has been observed among the growth traits for duck and mucin gene has been observed to play a role in growth characteristics. Hence mucin gene may be regarded as a promising gene affecting growth traits for indigenous duck, Bengal. This is the first report for role of mucin gene affecting growth traits.

Future study involves validation of these work with more number of ducks. Mucin gene may be regarded as an important gene affecting growth traits. Marker assisted selection or genomic selection posses the future scope of the study.Other similar genes may be studied for growth traits for a better conclusive result.

## Acknowledgement

The authors are thankful to Department of Science and Technology, Govt. of India (Grant no. EMR/2016/003554) for providing the financial support. The technical and financial support by Vice-Chancellor, West Bengal University of Animal and Fishery Sciences is duly acknowledged. Thanks to Director, AH & VS, Animal Resource Development Department, Govt. of West Bengal.

